# Ending transmission of SARS-CoV-2: sterilizing immunity using an intranasal subunit vaccine

**DOI:** 10.1101/2022.07.14.500068

**Authors:** Ankita Leekha, Arash Saeedi, Samiur Rahman Sefat, Monish Kumar, Melisa Martinez-Paniagua, Adrian Damian, Rohan Kulkarni, Ali Rezvan, Shalaleh Mosoumi, Xinli Liu, Laurence JN Cooper, Manu Sebastian, Brett Hurst, Navin Varadarajan

## Abstract

Immunization programs against SARS-CoV-2 with commercial intramuscular (IM) vaccines prevent disease but not infections. The continued evolution of variants of concern (VOC) like Delta and Omicron has increased infections even in countries with high vaccination coverage. This is due to commercial vaccines being unable to prevent viral infection in the upper airways and exclusively targeting the spike (S) protein that is subject to continuous evolution facilitating immune escape. Here we report a multi-antigen, intranasal vaccine, NanoSTING-NS that yields sterilizing immunity and leads to the rapid and complete elimination of viral loads in both the lungs and the nostrils upon viral challenge with SARS-CoV-2 VOC. We formulated vaccines with the S and nucleocapsid (N) proteins individually to demonstrate that immune responses against S are sufficient to prevent disease whereas combination immune responses against both proteins prevents viral replication in the nasal compartment. Studies with the highly infectious Omicron VOC showed that even in vaccine-naïve animals, a single dose of NanoSTING-NS significantly reduced transmission. These observations have two implications: (1) mucosal multi-antigen vaccines present a pathway to preventing transmission and ending the pandemic, and (2) an explanation for why hybrid immunity in humans is superior to vaccine-mediated immunity by current IM vaccines.

## INTRODUCTION

Humanity is in the middle of one of the largest vaccination campaigns to protect all people against the respiratory virus SARS-CoV-2 and coronavirus-induced disease (COVID-19). mRNA (*e.g*., BNT162b2 and mRNA-1273) and adenovirus vector vaccines (*e.g*., ChAdOx1 nCoV-19) have been delivered intramuscularly (IM) to billions of recipients^1,2^. The evolution of variants of concern (VOC) like the Omicron VOCs has caused a massive increase in infections even in countries with high vaccination coverage ^3^. This increased frequency of infections combined with laboratory data that supports increased infectivity and immune escape by the variants has seeded concerns that we will end up in the cumbersome perpetual cycle of immunization trying to keep pace with evolving variants ^4-7^.

There are two primary concerns with the commercial vaccines against SARS-CoV-2. First, while the IM route of administration elicits robust systemic immunity leading to the prevention of disease, they do not prevent viral infection in the upper airways. Unsurprisingly, the IM vaccines have demonstrated variable protection against upper-airway infection in preclinical models, with some offering no protection^8,9^. In humans, this led to both vaccinated and unvaccinated people harboring virus in the nostrils that facilitates transmission even by immunized individuals^10,11^. Moreover, the ability of the upper airways to serve as reservoirs facilitates viral evolution, and with the waning of vaccine-induced immunity over time, can enable the priming of new infections in vaccinated hosts^12,13^. The second concern is that the S protein dominates the vaccine landscape against SARS-CoV-2 as the immunogen^14^. Since the S protein is essential for viral entry into host cells, it serves as the preferred target for eliciting neutralizing antibodies^14,15^. Although correlates of vaccine-induced protection have not been established, there is strong evidence that neutralizing antibodies, and specific antibodies targeting the receptor-binding domain (RBD) of the S protein, are likely predictors of vaccine efficacy and disease prevention^16,17^. The S protein, however, by being on the surface of the virion, is under constant evolutionary pressure to escape the host immune system while preserving viral entry^18,19^. Unsurprisingly, as the virus has spread globally, variants less susceptible to antibodies elicited by vaccines have evolved, necessitating modified vaccine manufacturing and continued booster immunizations^19-21^.

Among other potential viral protein targets, the nucleocapsid (N) protein is expressed at elevated levels during infection and is highly immunogenic^22^. Studies tracking convalescent patient sera confirm robust antibody and T-cell responses against the N protein^23-26^. The primary function of the N protein is to package the viral genome into ribonucleoprotein complexes and to facilitate transcription while promoting escape from innate immunity (suppression of Th1 type I interferons, IFNs)^22^. Since the N protein performs multiple essential functions for the virus, it tends to accumulate fewer mutations resulting in the N protein of SARS-CoV-2 having 90 % homology to SARS-CoV^27^. These attributes make the N protein a candidate for vaccine-induced immunity^28^. Indeed, T-cell-dependent mechanisms can confer at least partial protection against the original Wuhan strain after IM vaccine candidates immunizing with the N protein^29^. However, preclinical studies have shown that transfer of anti-N immune sera failed to protect against SARS-CoV-2 infection in an adapted mouse model^30^ which is consistent with antibodies against the N protein not being neutralizing as this protein is unassociated with viral entry. Furthermore, intradermal vaccination with the SARS-CoV N protein worsened infection and pneumonia due to T helper 2 (Th2) cell-biased responses^31^. This concern of enhanced respiratory disease mediated by Th2 responses has shifted the focus away from the SARS-CoV-2 N protein-based vaccines despite the potential for protective T-cell responses.

Mucosal vaccines can stimulate robust systemic and mucosal immunity, but the quality and quantity of the immune response elicited upon mucosal vaccination depends on the appropriate adjuvant. We had previously reported that liposomally encapsulated endogenous STING agonist (STINGa, 2’-3’ cGAMP), termed NanoSTING, functions as an excellent mucosal adjuvant that elicits strong humoral and cellular immune responses upon intranasal vaccination^32^. Here we report that a multi-antigen intranasal subunit vaccine, NanoSTING-NS, delivers sterilizing immunity by eliminating the virus from the nose and lung. Our data provide a pathway to sterilizing immunity against even highly infectious variants and have important implications for understanding the quality of immunity elicited by natural infection compared to the current generation of vaccines targeting only the S protein.

## RESULTS

### Preparation and Characterization and NanoSTING-S vaccine

NanoSTING is a liposomal adjuvant that comprises pulmonary surfactant-biomimetic nanoparticle formulated STINGa and enables mucosal immunity (**Figure S1A**)^32,33^. We synthesized NanoSTING, and dynamic light scattering (DLS) showed that the mean particle diameter of NanoSTING was 137 nm, with a polydispersity index (PDI) of 24.5 % (**Figure S1B**) and a mean zeta potential of -63.5 mV (**Figure S1C**). We confirmed the ability of NanoSTING to induce IFN responses (IRF) using the THP-1 monocytic cells modified to conditionally secrete luciferase downstream of an IRF promoter. We stimulated THP-1 dual cells with NanoSTING and measured luciferase activity in the conditioned supernatant (**Figure S1D**) to show that secretion was maximal at 24 hours. We used recombinant trimeric S-protein to formulate the vaccine based on the SARS-CoV-2 B.1.351 (Beta VOC) as the immunogen^34^ (**Figure 1A**). SDS-PAGE under reducing conditions showed that the protein migrated between 180 and 250 kDa confirming extensive glycosylation (**Figure S2A**). Upon incubation with NanoSTING, the S protein was adsorbed onto the liposomes with NanoSTING-S displaying a mean particle diameter of 144 nm (PDI 25.9 %), and a mean zeta potential of −54.8 mV (**Figures S2B & S2C**). Unlike the trimeric S protein known to aggregate in solution, we tested NanoSTING-S after 9 months of storage at 4 °C. We found no evidence of aggregation, concluding that vaccine formulation is stable at 4 °C (**Figures S2D & S2E**).

**Figure 1:**
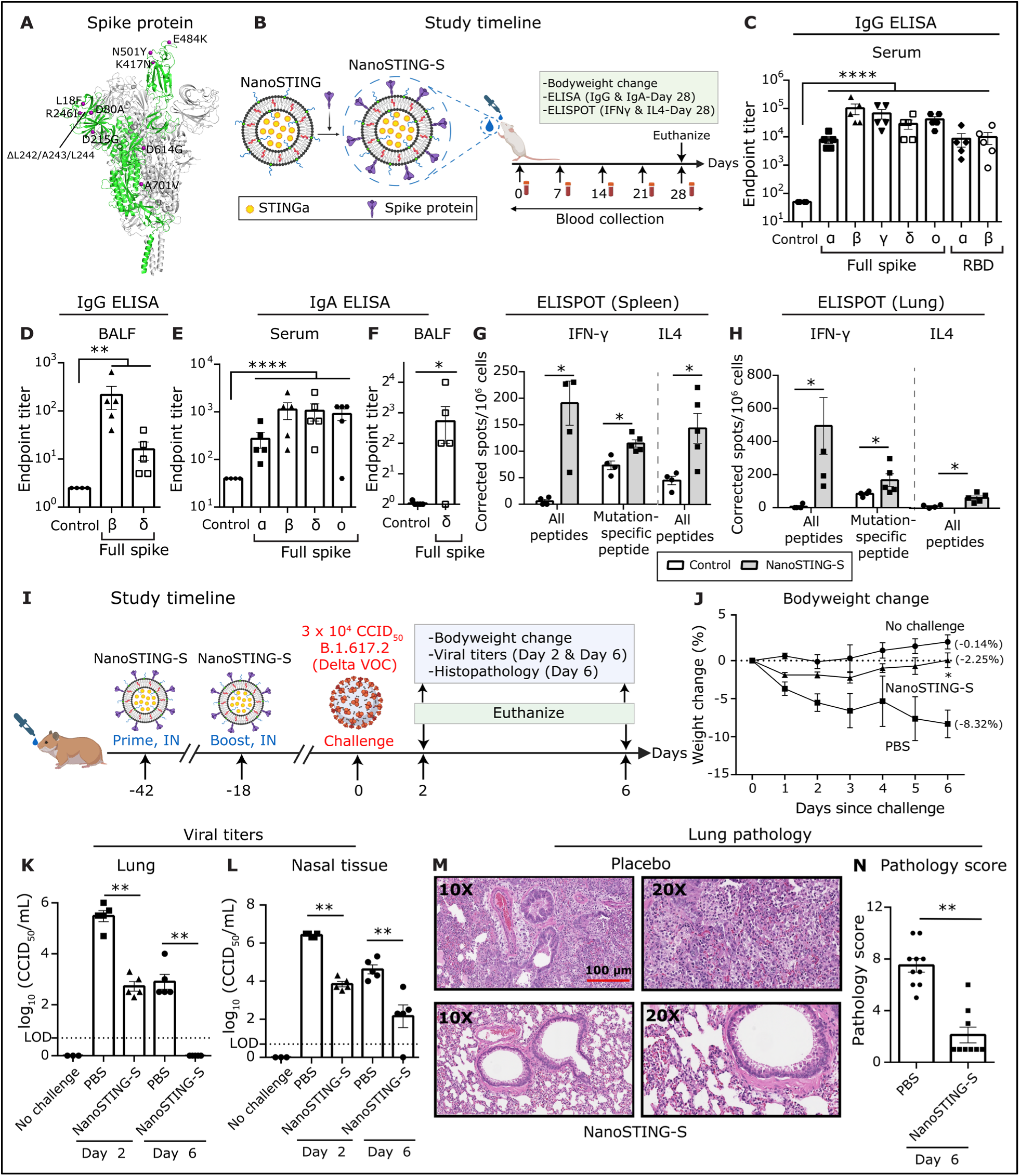
NanoSTING-S vaccine yields cross-reactive humoral and cellular immunity in mice and provides protective efficacy against Delta VOC in hamsters. **(A)** 3D structure of trimeric S protein (B.1.351) with the twelve mutations indicated (PDB: 7VX1)^77^. **(B)** Study timeline: We immunized Balb/c mice (n=5/group) with a single dose of NanoSTING-S intranasally (IN) followed by the collection of serum every week. We monitored the bodyweights of the animals every week after the immunization. We euthanized the animals at day 28 and then collected BALF, serum, lungs, and spleen. Bodyweight change, ELISA (IgG & IgA), and ELISPOT (IFNγ and IL4) were used as primary endpoints. Naïve Balb/c mice were used controls (n=4/group). **(C-F)** Humoral immune responses in the serum and BALF were evaluated using S-protein-based IgG & IgA ELISA. **(G, H)** Splenocytes (G) or lung cells (H) were stimulated *ex vivo* with overlapping peptide pools, and IFNγ & IL4 responses were detected using an ELISPOT assay. **(I)** Experimental set up for challenge study in hamsters: We immunized Syrian golden hamsters (n=10/group) intranasally with two doses of NanoSTING-S (first dose at day -42 and second dose at day -18, and challenged the hamsters intranasally with 3 × 10^4^ CCID_50_ of the SARS-CoV-2 Delta VOC on day 0. Post challenge, we monitored the animals for 6 days for changes in the bodyweight. We euthanized half of the hamsters on day 2 and other half at day 6 for histopathology of the lungs, with viral titers of lung and nasal tissues measured on day 2 and day 6. **(J)** Percent bodyweight change of hamster compared to the baseline at the indicated time intervals. **(K, L)** Viral titers measured by plaque assay in lungs and nasal tissues post day 2 and day 6 of infection. The dotted line indicates the limit of detection of the assay (LOD). **(M, N)** Pathology score and a representative hematoxylin and eosin (H&E) image of the lung showing histopathological changes in hamsters treated with NanoSTING-S and PBS; all images were acquired at 10x & 20×; scale bar, 100 µm. *For ELISA, ELISPOT, viral titers and lung histopathology data, analysis was performed using a Mann-Whitney test. Vertical bars show mean values with error bar representing SEM. Each dot represents an individual hamster or mouse. Weight data was compared via mixed-effects model for repeated measures analysis. Lines depict group mean bodyweight change from day 0; error bars represent SEM. Asterisks indicate significance compared to the placebo-treated animals at each time point*. *\*\*\*\*p < 0.0001; ***p < 0.001; **p < 0.01; *p < 0.05; ns: not significant.* *See also Figures S1, S2 & S3*

### Single-dose immunization of mice with NanoSTING-S vaccine yields cross-reactive humoral and cellular immunity against SARS-CoV-2

We immunized Balb/c mice with a single intranasal dose of NanoSTING-S (**Figure 1B**) and observed no clinical symptoms, including weight loss, during the entire 28-day period (**Figure S3**). We conducted ELISA on day 28 to quantify binding to both full-length S proteins and the RBDs, with the latter serving as a surrogate for neutralization. We observed robust serum IgG titers not only against Beta (B.1.351) but also against Alpha (B.1.1.7), Gamma (P.1), Delta (B.1.617.2), and Omicron (B.1.1.529) S proteins. We also observed high serum IgG titers against the RBDs from both the Beta and Alpha VOCs and high IgG titers against the full-length Beta and Delta spike proteins in bronchoalveolar lavage fluid (BALF, **Figure 1C-D**). As IgA-mediated protection is an essential component of mucosal immunity for respiratory pathogens, we confirmed the role of intranasal NanoSTING-S as a mucosal adjuvant. We detected elevated serum IgA responses against all spike protein variants tested, although BALF IgA titers against full-length delta spike protein were weaker (**Figure 1E & 1F**). At day 28, immunized mice showed robust and significant Th1/Tc1 responses by ELISPOT in both the spleen and the lung (**Figures 1G & 1H**). We stimulated the spleen and lung cells with a pool of peptides containing mutations (B.1.351) in the S protein that differs from the Wuhan S protein. We observed a significant Th1 response against these mutation-specific S peptides confirming a broad T-cell response that targets both the conserved regions and the mutated regions of the S protein (**Figures 1G & 1H**). In contrast to the Th1/Tc1 responses, the Th2 responses were weaker but detectable (**Figures 1G & 1H**). Collectively, these results established that NanoSTING acts as a mucosal adjuvant and that even a single-dose immunization with NanoSTING-S yielded robust IgG, IgA, and Th1/Tc1 responses that are cross-reactive against multiple VOCs.

### NanoSTING-S elicited immune responses confers protection against the Delta VOC

The Syrian golden hamster (*Mesocricetus auratus)* challenge model was used to assess the protective efficacy of NanoSTING-S. This animal model replicates COVID-19 severe disease in humans with infected animals demonstrating rapid weight loss, very high viral loads in the lungs, extensive lung pathology, and even features of long COVID^35,36^. Additionally, unlike the K18-hACE2 transgenic mouse model, hamsters recover from the disease and hence offer the opportunity to study the impact of treatments in the lungs (disease) and nasal passage (transmission)^35,37^. We chose the Delta VOC to infect the animals for two reasons: (1) this VOC causes severe lung damage, and (2) Delta-specific S-mutations, including L452R and T478K within the RBD are absent in our immunogen (**Figure 1A**) and provided an opportunity to assess cross-protection. We administered two doses of intranasal NanoSTING-S 24 days apart to hamsters which were subsequently challenged with the Delta VOC through the intranasal route (**Figure 1I**). Animals in the sham-vaccinated group showed a mean peak weight loss of 8.3 %. By contrast, animals vaccinated with NanoSTING-S were largely protected from weight loss (**Figure 1J**, mean peak weight loss of 2.3 %), consistent with the results obtained by adenovirally vectored IM vaccines challenged with either the Wuhan or Beta strains ^38^. We sacrificed half of the animals at day 2 (peak of viral replication) and the other half at day 6 (peak of weight loss in unimmunized animals) to quantify viral titers. NanoSTING-S reduced infectious viral loads in the lung by 300-fold by day 2 compared to sham-vaccinated animals, and by day 6, infectious virus was undetectable in all animals (**Figure 1K**). Viral replication in the lung of the animal models clinical human disease and death, while viral replication in the nasal compartment models human transmission. Immunization with NanoSTING-S reduced infectious viral loads in the nasal compartment by 380-fold by day 2 compared to unimmunized animals. By day 6, vaccinated animals showed a further significant reduction in the infectious virus (**Figure 1L**). To examine the pathobiology of viral infection, we analyzed the lung tissue on day 6 after the challenge using an integrated scoring rubric (range from 1-12) to quantify host immune response and disease severity. We recorded immune cell infiltration and widespread viral pneumonia in the lungs of sham-vaccinated hamsters, whereas vaccinated animals revealed minimal evidence of invasion by inflammatory cells or alveolar damage (**Figures 1M & 1N**). In aggregate, hamsters vaccinated with NanoSTING-S when challenged with the Delta VOC were protected in the lung against heterologous VOC and partially protected in the nasal passage. The reduction in viral loads in the nasal compartment suggests an advantage of mucosal vaccination to reduce transmission^39^.

### Modeling of the immune response against both S- and N-proteins predict synergistic protection

The results from the NanoSTING-S experiments demonstrated that the immune responses protect against disease in the lung but are insufficient to eliminate viral infection/replication in the nasal passage as a surrogate for transmission. A mathematical model was used to help understand if a multi-antigen vaccine comprising both S- and N-proteins (NanoSTING-NS) can offer improved protection^40^. We established the model to track the viral load in the nasal passage by fitting the parameters to reflect longitudinal viral titers from infected patients (**Figure 2A**). Vaccine-induced neutralizing antibody responses against the S-protein serve as *de novo* blockers of viral entry and impede viral production through immune effector mechanisms. We modeled a range (40 to 100 %) of vaccine efficacies (directed only against the S-protein) to account for the differences in protection, specifically in the nasal compartment, and investigated the influence on viral elimination. The model revealed a reduction in viral load between 35 % to 90 % when the S-vaccine efficacy in the nasal compartment varied from 40 to 80 % (**Figure 2B**). Anti-S vaccine efficacy >80% in the nasal compartment is difficult to accomplish even with mucosal immunization (some IM vaccines offer no significant nasal immunity) and can hence explain the inability to prevent nasal replication^8,9^. We next modeled a mucosal vaccine based exclusively on the N-protein. For the mucosal N-protein vaccine, we anchored to a mechanism of protection through induction of cytotoxic T cell responses that kill virally infected cells, thus reducing the number of cells capable of producing/propagating the virus. Under this scenario, the model predicted that the killing rate constant of cytotoxic T cells (CTL) would have to be 8.5 per day to achieve a 99% reduction in viral loads (**Figure 2C**). This value is at-least 10-fold higher than the predicted/measured killing capacity of CD8^+^ T cells *in vivo*, and hence it is not surprising that single-antigen N-based vaccines do not confer protection^41,42^. To quantify if multi-antigen vaccines can offer synergistic protection, we modeled combined protection by including S-directed vaccines that offer partial protection (primarily antibody-mediated) in the nasal compartment with the cytotoxic T cell responses against the N protein. We tested a range of S-protein vaccine efficacies (40 to 100 %) in the nasal compartment in combination with cytotoxic N responses (**Figure 2D**). The model predicted that a physiologically relevant CTL killing rate of 0.4 and 0.6 day per day would lead to a 1,000- and 10,000-fold reduction in peak viral load, respectively, when the efficacy of the spike vaccine was only 80% (**Figures 2D-red box & S4**). Indeed, studies in humans infected with COVID-19 have demonstrated a robust and long-lived CTL response in the nasal compartment and that CD8^+^ T cells specific for the N protein can directly inhibit viral replication^43,44^. Collectively, these results from modeling predicted that combination vaccines targeting S and N proteins can mediate synergistic protection in eliminating viral replication in the nasal compartment.

**Figure 2:**
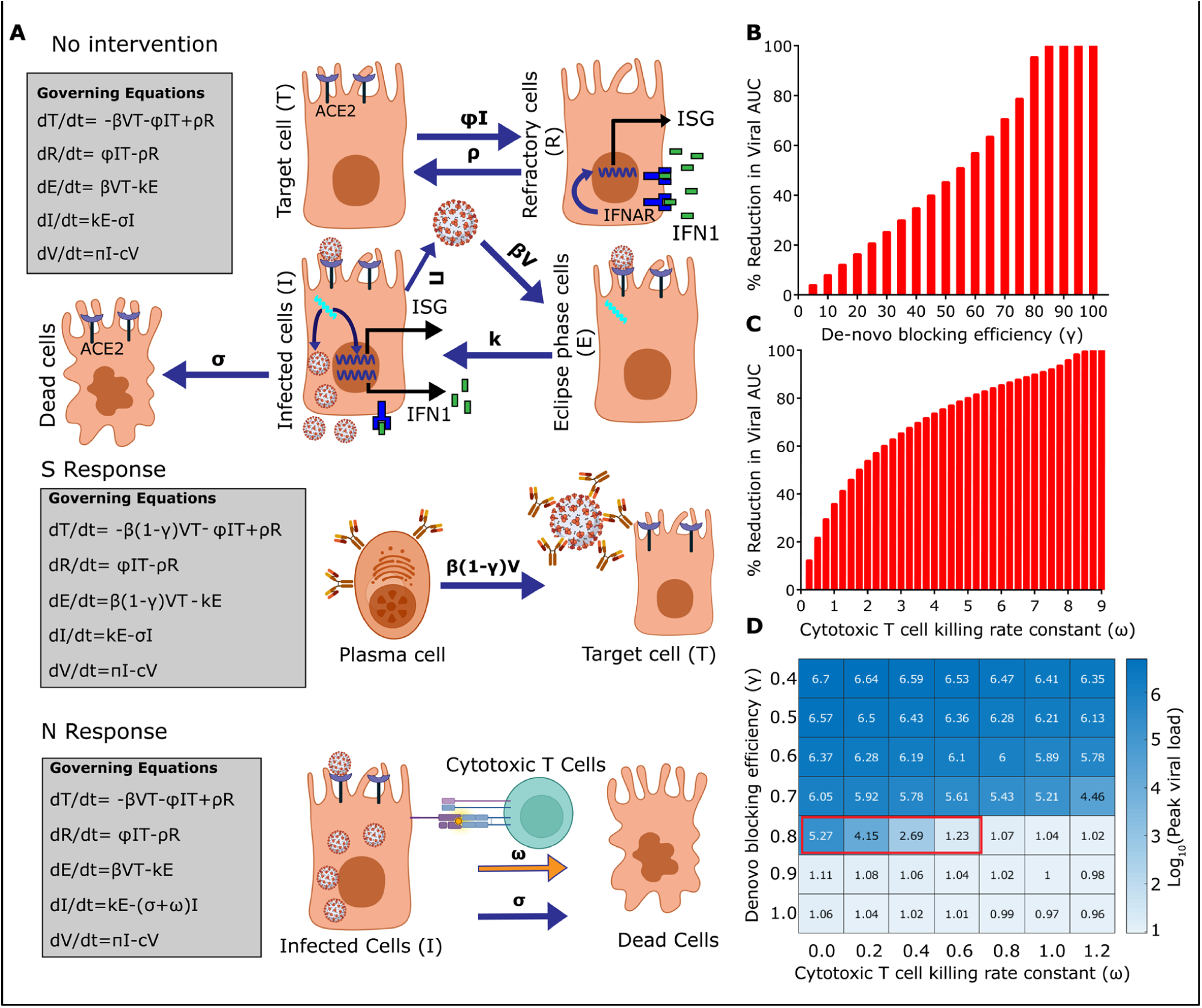
Quantitative modeling of the combined immune response against both proteins predict synergistic protection. **(A)** Schematic and governing equations describing viral dynamics without vaccination, with spike protein immunization, or nucleocapsid protein immunization (IFNAR: interferon-α/β receptor, IFN1: type-I interferons, ISG: interferon-stimulated gene). **(B)** Percent reduction in viral AUC with increasing de-novo blocking efficiency (antibodies against the spike protein). **(C)** Percent reduction in viral AUC upon cytotoxic T cell-mediated killing of infected cells. **(D)** Heatmap showing the effectiveness of combined effect of de-novo blocking (S response) and T cell-mediated killing (N response). The red box indicates the synergistic effect of N and S response in achieving sterilizing immunity. *See also Figure S4*

### Single-dose immunization of mice with NanoSTING-NS vaccine yields balanced humoral and cellular immunity and sterilizing immunity

We formulated vaccines containing both antigens to test the model that the immune response against both the S and N proteins can be synergistic (**Figure 3A**). We initially performed immunogenicity experiments in mice with 10 µg each of the recombinant N and S proteins adjuvanted with NanoSTING. We observed that while 100% of animals seroconverted and showed IgG responses against the S protein, seroconversion against the N protein was variable (40-80 %) [not shown]. We accordingly modified the mass ratio of N:S protein (2:1) and adjuvanted it with NanoSTING to formulate NanoSTING-NS (**Figure 3A**). The NanoSTING-NS displayed a mean particle diameter of 142 nm (PDI 26.2 %) and a mean zeta potential of −48.4 mV (**Figures S5A & 5B**). We tested NanoSTING-S after 9 months of storage at 4 °C, and confirmed that it displayed excellent stability, like the NanoSTING-S vaccine (**Figures S5C & S5D**). Single-dose intranasal vaccination in mice with NanoSTING-NS was safe (**Figure S6**) and yielded robust serum IgG titers against the N protein and full-length S protein variants at day 35 (**Figure 3B**). We documented robust antigen-specific, cross-reactive IgG responses in the BALF (**Figure 3C**) and observed cross-reactive IgA responses in the serum and BALF at day 35 (**Figures 3D & 3E**). We measured Th1 and Th2 responses against the N- and S-proteins in both the spleen and the lung at day 51 (**Figures 3F & 3G**) and observed no significant Th2 response (IL4) in both tissues. Based on these promising immunogenicity data in mice, we evaluated the protective efficacy of NanoSTING-NS in hamsters. We vaccinated hamsters intranasally with two doses of NanoSTING-NS and challenged the immunized hamsters with the Delta VOC through the intranasal route (**Figure 3H**). Animals immunized with NanoSTING-NS were completely protected from weight loss (mean peak weight loss of 0.8 %) [**Figure 3I**]. Like the results of the NanoSTING-S vaccine, NanoSTING-NS eliminated viral replication in the lung by day 6 post-challenge (**Figure 3J**), suggesting that S-specific immune responses are the dominant factor in providing immunity in the lung. In the nasal compartment, NanoSTING-NS showed a significant reduction in infectious viral particles by day 2 even in comparison to NanoSTING-S, and significantly, by day 6 there was a complete elimination of infectious viral particles in the nasal tissue of the NanoSTING-NS vaccinated animals (**Figure 3K**). Pathology also confirmed that vaccinated and challenged animals had minimal evidence of invasion by inflammatory cells or alveolar damage (**Figures 3L & 3M**). In aggregate, these results illustrate that NanoSTING-NS can provide sterilizing immunity.

**Figure 3:**
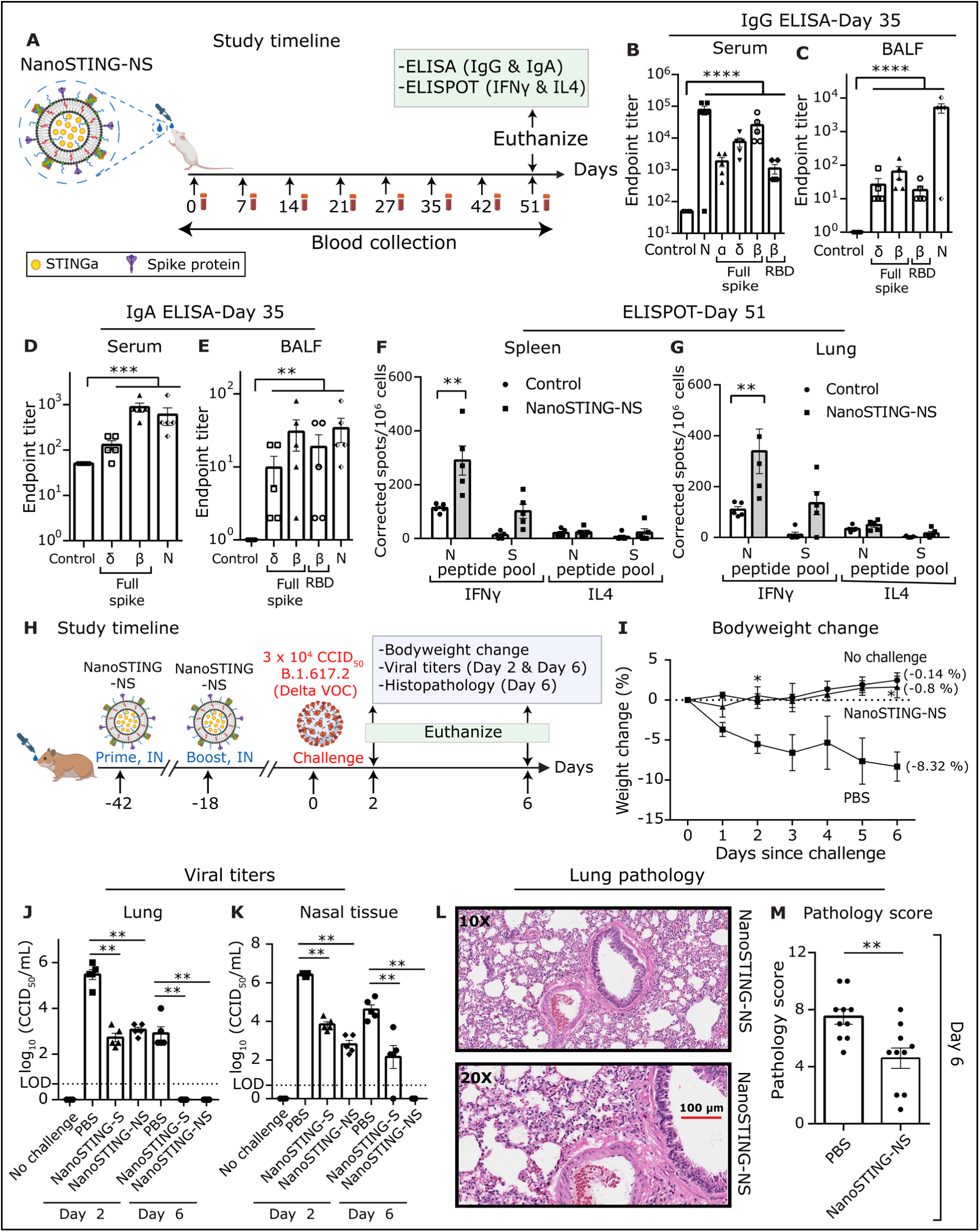
NanoSTING-NS vaccine yields balanced humoral and cellular immunity targeting both proteins and yields sterilizing immunity. **(A)** Experimental set up: We immunized two groups (n=5/group) of mice by intranasal administration with NanoSTING-NS followed by the collection of serum every week. We monitored bodyweights of the animals every week after the immunization until the end of the study. We euthanized the animals at day 51 followed by the collection of BALF, serum, lungs, and spleen. Bodyweight change, ELISA (IgG & IgA), and ELISPOT (IFNγ and IL4) were used as primary endpoints. Naïve Balb/c mice were used controls (n=5/group). **(B-E)** Humoral immune responses in the serum and BALF were evaluated using S-protein based IgG & IgA ELISA. **(F, G)** Splenocytes (F) or lung cells (G) were stimulated *ex vivo* with overlapping peptide pools, and IFNγ & IL4 responses were detected using an ELISPOT assay. **(H)** Timeline for challenge study done in Syrian golden hamsters: We immunized hamsters intranasally with two doses of NanoSTING-NS (first dose at day -42 and the second dose at day -18) and challenged the hamsters intranasally with 3 × 10^4^ CCID_50_ of the SARS-CoV-2 Delta VOC on day 0. Post challenge, we monitored the animals for 6 days for changes in the bodyweight. We euthanized half of the hamsters on day 2 and the other half at day 6 for histopathology of the lungs, with viral titers of lung and nasal tissues measured on day 2 and day 6. **(I)** Percent bodyweight change of hamster compared to the baseline at the indicated time intervals. **(J, K)** Viral titers measured by plaque assay in lungs and nasal tissues post day 2 and day 6 of infection. The dotted line indicates LOD. **(L)** Pathology score and a representative hematoxylin and eosin (H&E) image of the lung showing histopathological changes in hamsters treated with NanoSTING-NS and PBS; all images were acquired at 10x & 20×; scale bar, 100 µm. *For ELISA, ELISPOT, viral titers and lung histopathology data, analysis was performed using a Mann-Whitney test. Vertical bars show mean values with error bar representing SEM. Each dot represents an individual hamster or mouse. Weight data was compared via mixed-effects model for repeated measures analysis. Lines depict group mean bodyweight change from day 0; error bars represent SEM. Asterisks indicate significance compared to the placebo-treated animals at each time point*. *\*\*\*\*p < 0.0001; ***p < 0.001; **p < 0.01; *p < 0.05; ns: not significant.* *See also Figures S5 & S6*

### Immunization of mice with NanoSTING-N yields durable humoral and cellular immunity but is not sufficient to confer protection against Delta VOC

To quantify the role of anti-N immunity in mucosal protection, we characterized the immune response elicited against the N protein by formulating NanoSTING-N and testing it in mice. Independent studies with K18-hACE2 mice immunized with viral vector-based N protein and challenged the early lineage variants (Wuhan and Alpha VOC) revealed mixed results with either partial or a complete lack of protection^29,45^. The predicted structure of the SARS-CoV-2 N protein comprises an RNA binding domain, a C-terminal dimerization domain, and three intrinsically disordered domains that promote phase separation with nucleic acids^46^ (**Figure S7A**). We confirmed the functional activity of the protein by assaying binding to plasmid DNA based on the quenching of the fluorescent DNA condensation probe DiYO-1 (**Figure S7C)**^47^. To formulate the vaccine, NanoSTING-N, we mixed the N protein with NanoSTING to allow the adsorption of the protein onto the liposomes (**Figure 4A**). The formulated NanoSTING-N had a mean particle diameter of 107 nm (PDI 20.6 %), and zeta potential of -51 mV (**Figures S7E & S7F**). Although the recombinant N protein showed a strong propensity for aggregation upon storage at 4 °C for 6 months, NanoSTING-N was stable with no change in size or zeta potential (**Figures S7G & S7H**). Consistent with our NanoSTING-NS studies, we immunized two groups of mice by intranasal administration with either 10 µg (NanoSTING-N10) or 20 µg of N protein (NanoSTING-N20, **Figures 4A, and S8**). The serum IgG responses at both doses were similar at day27, although the IgG titers elicited by the NanoSTING-N20 were higher than NanoSTING-N10, the difference was not significant (**Figure 4B**).

**Figure 4:**
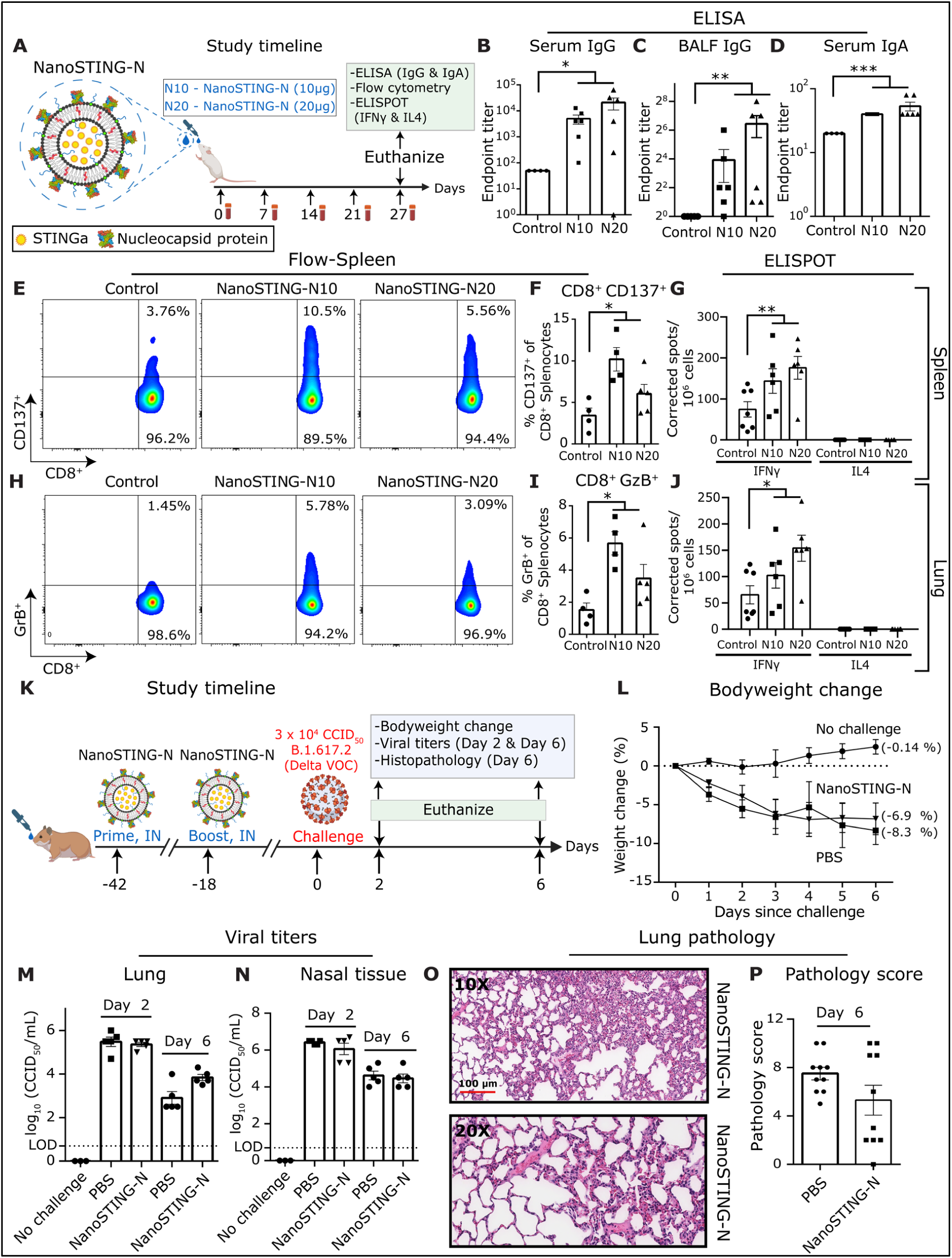
NanoSTING-N vaccine yields durable humoral and cellular immunity in mice but is insufficient to confer protection against the highly infectious Delta VOC in hamsters. **(A)** Experimental set up: We immunized two groups (n=5-6/group) of mice by intranasal administration with NanoSTING-N10 or NanoSTING-N20 followed by serum collection every week. We monitored the bodyweights of the animals every week after the immunization until the end of the study. We euthanized the animals at day 27 and then collected BALF, serum, lungs, and spleen. Bodyweight change, ELISA (IgG & IgA), flow cytometry (CD8^+^ T cells), and ELISPOT (IFNγ and IL4) were used as primary endpoints. Naïve Balb/c mice were used controls (n=4/group). **(B, C)** Humoral immune responses in the serum and BALF were evaluated using N-protein based IgG ELISA. **(D)** Humoral immune responses in the serum were evaluated using N-protein-based IgA ELISA. **(E-G)** Splenic CD8^+^ T cells were stimulated *ex vivo* with overlapping peptide pools, and (E, F) CD137 expression was quantified by flow cytometry (G) IFNγ & IL4 responses were detected using an ELISPOT assay **(H, I)** Splenic CD8^+^ T cells were stimulated *ex vivo* with overlapping peptide pools, and (H, I) GzB expression was quantified by flow cytometry **(J)** IFNγ & IL4 ESLIPOT from lung cells stimulated *ex vivo* with indicated peptide pools. **(K)** Experimental set up for challenge studies in Syrian golden hamsters. We immunized hamsters (n=10/group) intranasally with two doses of NanoSTING-N (first dose at day -42 and the second dose at day -18, and challenged the hamsters intranasally with 3 × 10^4^ CCID_50_ of the SARS-CoV2 Delta VOC on day 0. Post challenge, we monitored the animals for 6 days for changes in bodyweight. We euthanized half of the hamsters on day 2 and other half at day 6 for histopathology of the lungs, with viral titers of lung and nasal tissues measured on day 2 and day 6. **(L)** Percent bodyweight change of hamster compared to the baseline at the indicated time intervals. **(M, N)** Viral titers measured by plaque assay in lungs and nasal tissues post day 2 and day 6 of infection. The dotted line indicates LOD. **(O, P)** Pathology score and a representative H&E image of the lung showing histopathological changes in hamsters treated with NanoSTING-N and PBS; all images were acquired at 10x & 20×; scale bar, 100 µm. *For ELISA, flow cytometry, ELISPOT, viral titers and lung histopathology data, analysis was performed using a Mann-Whitney test. Vertical bars show mean values with error bar representing SEM. Each dot represents an individual hamster or mouse. Weight data was compared via mixed-effects model for repeated measures analysis. Lines depict group mean bodyweight change from day 0; error bars represent SEM. Asterisks indicate significance compared to the placebo-treated animals at each time point*. *\*\*\*\*p < 0.0001; ***p < 0.001; **p < 0.01; *p < 0.05; ns: not significant.* *See also Figures S7-S11*

In contrast to vaccination with the trimeric NanoSTING-S (early response at day 7), the kinetics of IgG responses were delayed, and responses were only observed at day 14 (**Figure S9**). Both, NanoSTING-N10 and NanoSTING-N20 yielded antigen-specific IgG responses in the BALF and IgA response in serum (**Figures 4C & 4D**). We examined the activation and function of N-protein-specific memory CD8^+^ T cells in the lungs and spleen using granzyme B (GzB) and the activation-induced marker CD137 (**Figure S10A**). Restimulation *ex vivo* with a pool of overlapping peptides derived from the N protein resulted in a significant increase in the frequency of activated (CD8^+^CD137^+^), and cytotoxic (CD8^+^GzB^+^) T cells in the spleen (**Figures 4E, 4F, 4H & 4I**) and to a lesser extent in the lung, of both the NanoSTING-N10 and NanoSTING-N20 vaccinated mice (**Figures S10B & S10C**). The overall frequencies of the lung resident CD8^+^CD103^+^ and CD103^+^CD69^+^CD8^+^ T cells were no different between the immunized animals and the control group (**Figures S10D & S10E**). NanoSTING-N10 and NanoSTING-N20 immunized mice showed robust and significant splenic and lung Th1/Tc1 responses (**Figures 4G & 4J**). We did not observe a measurable IL4 (Th2) response upon immunization with NanoSTING-N10 and NanoSTING-N20 (**Figures 4G & 4J**). Collectively, these results established that intranasal vaccination elicited strong Th1/Tc1 responses with no evidence of Th2 responses. To test the durability of the response, we immunized mice with NanoSTING-N20 and monitored the animals for 62 days (**Figure S11A**). NanoSTING-N20 vaccinated animals reported no weight loss (**Figure S11B**) and revealed robust serum IgG and IgA titers at day 62 (**Figure S11C & S11D)**. We also confirmed that the N-reactive Th1 responses were conserved in the spleen and lung at day 62 (**Figure S11E & S11F**). These results can be summarized as immunization with NanoSTING-N results in IgG and IgA immune responses and long-lived Th1/Tc1 but not deleterious Th2 immune responses.

Based on the immunogenicity data in mice, we evaluated the protective efficacy of NanoSTING-N in hamsters. We intranasally vaccinated hamsters with two doses of NanoSTING-N and challenged the immunized hamsters with the Delta VOC through the intranasal route (**Figure 4K**). Animals in both the vaccinated and sham-vaccinated groups showed significant weight loss (**Figure 4L)**. Consistent with the lack of protection from weight loss, infectious viral titers were no different in the lung or nasal passage on either day 2 or day 6 in both vaccinated and sham-vaccinated animals (**Figures 4M & 4N**). In addition, we observed that the aggregate pathology score of NanoSTING-N treated hamsters was not significantly different from sham-vaccinated animals, although the distribution of pathology scores appeared bimodal (**Figures 4O & 4P**). Collectively, the immunization and the challenge data are aligned with our mathematical model and illustrate that while NanoSTING-N elicits strong Tc1 responses, these responses are insufficient to prevent viral expansion in the absence of S-directed immunity.

### A single dose of NanoSTING-NS significantly reduces direct transmission of SARS-CoV-2 Omicron VOC

The current Omicron variants are transmitted very efficiently, and we next wanted to directly investigate if single immunization with NanoSTING-NS can mitigate the transmission of highly infectious VOC. We established a transmission experiment using the Omicron VOC (B.1.1.529) and two groups of Syrian golden hamsters. For group 1, we immunized the index hamsters with a single dose of NanoSTING-NS, and three weeks later, we infected with index hamsters with SARS-CoV-2 Omicron VOC. One day after infection, each index hamster was paired with a co-housed contact (unimmunized) hamster for 4 days, permitting transmission with direct contact (**Figure 5A**). We quantified the viral titers in the index and contact hamsters. As with the other strains of SARS-CoV-2 that we tested, immunization with NanoSTING-NS significantly reduced the viral titers in the nasal tissue of contact hamsters as compared to sham-vaccinated controls (**Figure 5B**). The index hamsters also showed reduced viral titers in nasal tissue at day 5 (**Figure 5C**). These results demonstrate that even a single dose of NanoSTING-NS is highly effective at mitigating the transmission of the Omicron VOC, which has implications for controlling the outbreak of respiratory pathogens.

**Figure 5:**
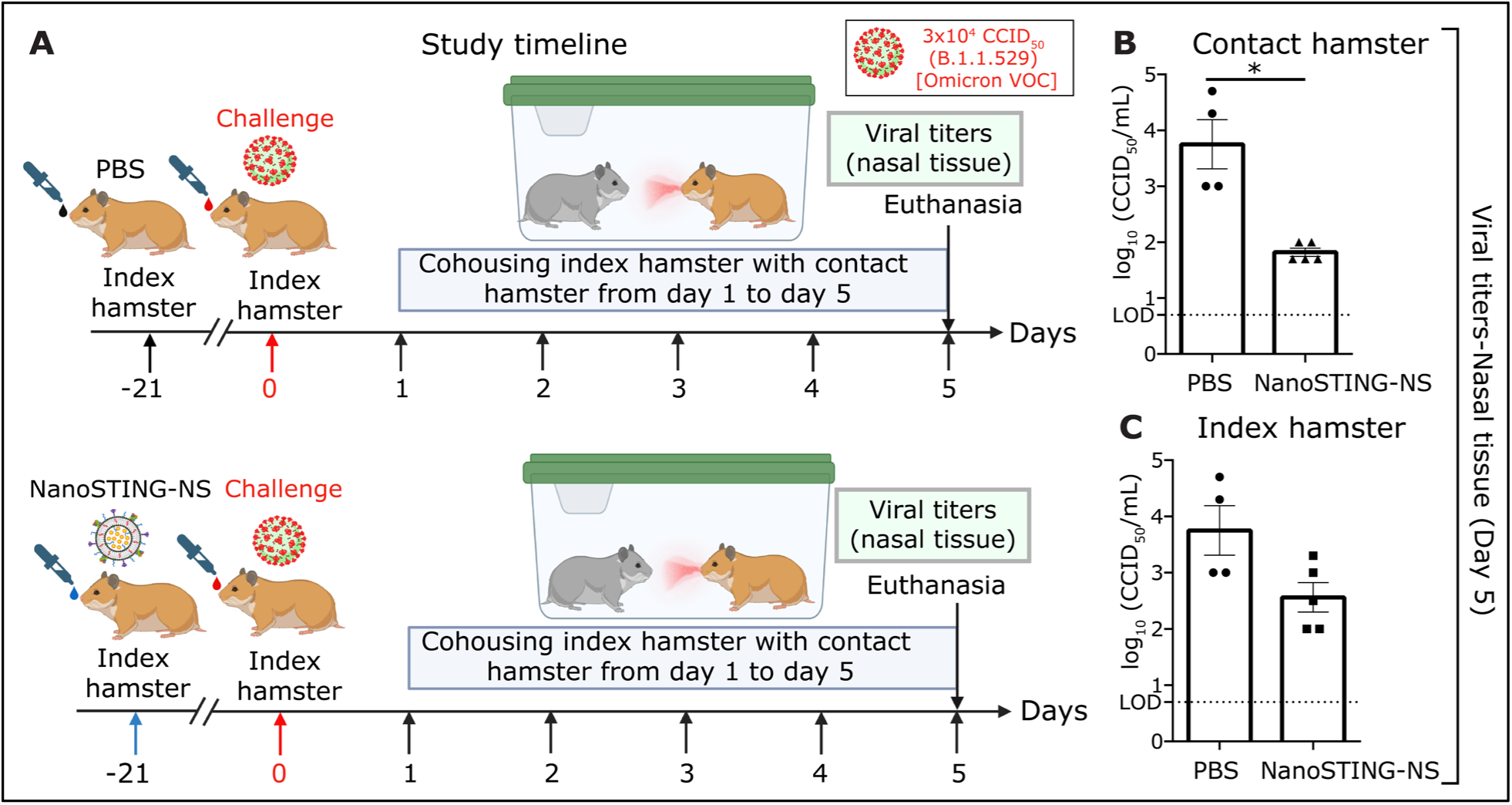
Intranasal administration of NanoSTING-NS limits transmission and viral replication of SARS-CoV-2 Omicron (B.1.1.529) VOC in hamsters. **(A)** Experimental set up: We immunized hamsters with a single dose of the intranasal NanoSTING-NS vaccine (n=5/group) or PBS (n=4/group) three weeks prior to infection with ∼3 × 10^4^ CCID_50_ of SARS-CoV-2 Omicron VOC (B.1.1.529). One day after the viral challenge, we co-housed the index hamsters in pairs with contact hamsters for 4 days in clean cages. We euthanized the contact and index hamsters on day 4 of cohousing. Viral titers in the nasal tissue of the index and contact hamsters were used as primary endpoints. **(B, C)** Infectious viral particles in the nasal tissue of contact and index hamsters at day 5 after viral administration post-infection were measured by plaque assay. The dotted line indicates LOD. *For viral titers, analysis was performed using a Mann-Whitney test. Vertical bars show mean values with error bar representing SEM. Each dot represents an individual hamster. Asterisks indicate significance compared to the placebo-treated animals*. *\*\*\*\*p < 0.0001; ***p < 0.001; **p < 0.01; *p < 0.05; ns: not significant*.

## DISCUSSION

Our study makes an essential contribution to achieving sterilizing immunity for SARS-CoV-2. We demonstrate that the NanoSTING-NS intranasal vaccine candidate NanoSTING-NS can eliminate viral loads both in the lung and in the upper airways of hamsters challenged with the Delta VOC. In direct transmission models, even a single dose of our vaccine was sufficient to significantly reduce transmission of the highly transmissible and currently dominant Omicron VOC in vaccine-naïve animals. An alternative intranasal vaccine candidate based on a viral vector expressing the S protein (ChAd-SARS-CoV-2-S) showed results similar to NanoSTING-S, demonstrating reduction but also variability in viral loads in the upper airways of K18-hACE2 mice challenged with chimeric viruses with spike genes corresponding to SARS-CoV-2 VOC (B.1.351 and B.1.1.28)^48^. It is predictable that the intranasal N protein-based vaccine (NanoSTING-N) is highly immunogenic but failed to protect the Delta VOC. Investigations with K18-hACE2 mice immunized with viral vector-based N protein and challenged with the early lineage variants (Wuhan and Alpha) show either a partial or complete lack of protection, similar to our results^29,45^. Remarkably, by combining the N- and S-proteins in the intranasal vaccine, NanoSTING-NS eliminated the virus not only in the lung (like NanoSTING-S) but also in the upper airways.

We propose a model based on our mice and hamster data (mechanistic studies are difficult to accomplish in hamsters due to the unavailability of appropriate hamster-specific reagents, and challenge models in K18-hACE2 mice are lethal with limited opportunity to study transmission). In the lung, the S-specific Ig (IgG and IgA) is sufficient to neutralize the virus. However, only IgA is present in the nasal cavity, which is insufficient to prevent viral entry into cells. In this context, the N-specific T-cell responses are hypothesized to be responsible for eliminating the residual virally infected cells, thereby achieving sterilizing immunity. An outcome of sterilizing immunity is reducing incidence of viral transmission between the infected and the uninfected, which we successfully demonstrated. The ability to elicit mucosal immune responses and directly dampen transmission even with the Omicron VOC differentiates NanoSTING-NS from IM multi-antigen vaccines^49,50^.

The ability to elicit multifactorial immunity in the nasal cavity has several implications for SARS-CoV-2 vaccines. First, since respiratory viruses like SARS-CoV-2 can access the brain through the olfactory mucosa in the nasal cavity, immunity in this compartment can prevent viral seeding to the brain. This, in turn, can prevent the entire spectrum of neurological complications ranging from the immediate loss of smell and taste to long-term complications like stroke^51,52^. Second, eliminating the virus in the nasal cavity of vaccinated recipients reduces the chance of viral evolution leading to breakthrough disease, especially in the context of waning immunity^53^. Allowing the virus to persist is a risky experiment in viral evolution with likely tragic consequences^54^. Eradicating the SARS-CoV-2 viral reservoir in humans provides the only reasonable path to moving past the pandemic and the perpetual cycle of repeated booster vaccinations. History provides a powerful example of the importance of vaccinating to prevent infections, not just disease, and sets a clinical precedent. Similar to the current COVID-19 commercial vaccines, the first inactivated polio vaccine in 1955 successfully prevented disease but not infection. The availability of the oral polio vaccine starting in 1960 paved the way for eliminating infection and eradicating polio. The availability of multi-antigen mucosal vaccines provides a pathway for humanity to move past SARS-CoV-2 outbreaks.

Systematically quantifying the immunity against multiple proteins also has important implications for understanding the nature of the immune response elicited upon infection versus vaccination targeting single proteins (all approved COVID-19 vaccines). Preclinical studies with hamsters show that intranasal challenge with the ancestral Wuhan strains results in 100% seroconversion and protects from subsequent reinfection by the Delta VOC^55^. Several observational studies profiling millions of humans support that hybrid immunity (one or more doses of vaccines and infection) is superior to two or even three doses of vaccines in preventing severe infection/hospitalization with emerging variants^56-58^. The superiority of hybrid immunity has been documented when the primary infection was with the Delta VOC, and now, more recently, with the Omicron VOC^59-62^. Mechanistic studies have uncovered important observations about the quality/quantity of the immune responses that explain the superiority of hybrid immunity over vaccine-induced immunity: (1) potent antibody responses with increased coverage that can neutralize multiple variants^63,64^, (2) the ability to elicit superior Fc-dependent functions^65^, and (3) unique subsets of T cells with the secretion of IFNγ and IL10^66^. Unfortunately, most mechanistic studies have focused on the immune response against the S protein, likely because the approved IM vaccines only target this protein.

Human studies have routinely documented that infection provides reliable anti-N protein antibody and T cell responses^67-69^. As our results with quantitative modeling and the NanoSTING-N vaccine illustrate, immunity solely against the N protein in the absence of robust antibody responses against the S protein is not protective against the SARS-CoV-2 viral challenge. The disadvantage of the immunity elicited by natural infections in humans is that the antibody response against the S protein is variable and depends on the type of SARS-CoV-2 variant and the severity of infection^70^. By contrast, vaccine-induced responses against the S protein are much more reliable and uniform. Multiple doses of vaccines ensure strong immunity (both antibodies and T cell responses) against the S protein. Our data with NanoSTING-S and data from multiple immunization studies illustrate that robust immunity against the S protein prevents severe disease. Hybrid immunity is thus able to derive the multifactorial benefit of responses in different anatomical compartments (mucosal vs systemic), against multiple antigens, and potentially against multiple variants (if infection was with any strain besides the ancestral strain). Evidence of the importance of this multifactorial immunity is available in human studies that document T-cell responses against multiple antigens and preclinical studies that have used multi-antigen vaccines^45,71,72^. With our data, we submit that the immunity elicited by NanoSTING-NS (multi-antigen) models hybrid immunity: it ensures mucosal immune responses against the N protein while preserving the strong systemic responses against the S protein.

In summary, we have developed and validated a multi-component intranasal NanoSTING-NS subunit vaccine candidate that eliminates SARS-CoV-2 in the upper airways, preventing transmission and provides a route for sterilizing immunity in humans.

## MATERIAL & METHODS

### Preparation of NanoSTING, NanoSTING-S, NanoSTING-N, and NanoSTING-NS

The liposomes contained DPPC, DPPG, Cholesterol (Chol), and DPPE-PEG2000 (Avanti Polar lipids) in a molar ratio of 10:1:1:1. To prepare the liposomes, we mixed the lipids in CH_3_OH and CHCl_3_, and used a vacuum rotary evaporator to evaporate them at 45°C. We dried the resulting lipid thin film in a hood to remove any residual organic solvent. Next, we added pre-warmed cGAMP (Chemietek) solution (3 mg/mL in PBS buffer at pH 7.4) to hydrate the lipid film. We mixed the hydrated lipids for an additional 30 min at an elevated temperature of 65 °C and subjected them to freeze-thaw cycles. Using a Branson Sonicator (40 kHz), we next sonicated the mixture for 60 min and used Amicon Ultrafiltration units (MW cut off 10 kDa) to remove the free untrapped cGAMP. Finally, we used PBS buffer to wash the NanoSTING (liposomally encapsulated STINGa) three times. We measured the cGAMP concentration in the filtrates against a calibration curve of cGAMP at 260 nm using Take3 Micro-Volume absorbance analyzer of Cytation 5 (BioTek). We calculated the final concentration of cGAMP in NanoSTING and encapsulation efficiency by subtracting the concentration of free drug in the filtrate.

To prepare NanoSTING adjuvanted subunit protein vaccine, we used a simple “mix and adsorb” approach. Briefly, (i) NanoSTING-S vaccine was prepared by gently mixing 10µg of trimeric spike protein-B.1.351 (Acrobiosystems, #SPN-C52Hk) with 20µg of NanoSTING. (ii) NanoSTING-N (Wuhan) (BEI, # NR-53797): Two different concentrations of the Nucleocapsid protein were taken: NanoSTING-N10 (10 µg of N protein) and NanoSTING-N20 (20 µg of N protein) were mixed separately with 20 µg of the NanoSTING. (iii) NanoSTING-N: 20µg of nucleocapsid protein-B.1.17 (Acrobiosystems, #NUN-C52H8) was mixed with 20µg of NanoSTING. (iv) NanoSTING-NS: 10µg of trimeric spike protein-B.1.351 (Acrobiosystems, #SPN-C52Hk) and 20µg of nucleocapsid protein-B.1.17 (Acrobiosystems, #NUN-C52H8) were mixed with 20µg of NanoSTING. All the vaccines were left on ice for a minimum of 1 h with constant slow shaking on the rocker.

### Stability studies for the formulated vaccines

We stored the NanoSTING, NanoSTING-S, NanoSTING-N, and NanoSTING-NS at 4°C for 6-9 months to check their stability. We measured the average hydrodynamic diameter and zeta potential of NanoSTING and all vaccine formulations using DLS and zeta sizer on Litesizer 500 (Anton Paar).

### Cell lines

THP-1 dual™ cells (NF-κB-SEAP IRF-Luc Reporter Monocytes) [InvivoGen, SanDiego, CA, thpd-nfis] was cultured in a humidified incubator at 37 °C and 5% CO_2_ and grown in RPMI 1640, 2 mM L-glutamine, 25 mM HEPES, 10% heat-inactivated fetal bovine serum (30 min at 56°C), 100 μg/ml Normocin™, Pen-Strep (100 U/ml-100 μg/ml). THP-1 Dual cells were grown in the presence of respective selection agents [100 mg/mL zeocin (InvivoGen, #ant-zn-1) and 10 mg/mL blasticidin (InvivoGen, #ant-bl-1)] every other passage to maintain positive selection of reporters.

### Cell stimulation experiments with luciferase reporter enzyme detection

We performed the THP-1 cell stimulation experiments using the manufacturer’s instructions (InvivoGen, CA, USA). First, we seeded the cells in 96 well plate at 1 × 10^5^ cells/well in 180 μL growth media. We then incubated the cells with 5 µg of NanoSTING at 37 °C for 24 h. To detect IRF activity, we collected 10 μL of culture supernatant/well at 12 h and 24 h and transferred it to a white (opaque) 96 well plate. Next, we read the plate on Cytation 7 (Cytation 7, Bio-Tek Instruments, Inc.) after adding 50 μL QUANTI-Luc™ (InvivoGen) substrate solution per well, followed by immediate luminescence measurement, which was given as relative light units (RLU).

### DNA Binding Assay

We performed the DNA binding studies as previously published^47^. To check the applicability of the assay for detecting DNA condensation, we used branched-chain PEI (Polyethylenimine) as a positive control (Sigma Chemical Co., St. Louis, MO # 408727). DiYO-1 (AAT Biorequest #17579) and plasmid (pMB75.6)-DNA were mixed in equal volumes (in 20 mM HEPES, 100 mM NaCl, pH = 7.4) to achieve a final concentration of 400 nM DNA phosphate and 8 nM DiYO-1, respectively. The solution was left at RT for 5 h before use. Next, we added PEI at different concentrations (R = 0, 1, 2, 5 where R is the molar ratio of PEI Nitrogen to DNA phosphate) to DNA-DiYO-1 solution, mixed for 1 min, and left for 2 h to equilibrate. We measured the fluorescence intensity of the solution at excitation and emission wavelength of 470 nm and 510 nm, respectively. We repeated the same procedure with SARS-CoV-2 N protein instead of PEI. To the DNA-DiYO-1 solution, we added the N protein at concentrations of 0.1 µM and 0.5 µM.

### Mice and immunization

All the animal experiments were reviewed and approved by UH IACUC. We purchased the female, 7–9-week-old BALB/c mice from Charles River Laboratories. Before immunization, we anesthetized the groups of mice (n=4-6/group) by intraperitoneal injection of ketamine (80µg/g of body weight) and xylazine (6µg/g of body weight). Then, we immunized the animals with (i) NanoSTING-S vaccine (ii) NanoSTING-N10 and NanoSTING-N20 (iii) NanoSTING-N (iv) NanoSTING-NS. All vaccines were freshly prepared.

### Bodyweight monitoring and sample collection

We monitored the bodyweight of the animals every 7 days until the end of the study after immunization. In addition, we collected the sera every week post-vaccination to detect the humoral immune response. We kept the blood at 25 °C for 10 min to facilitate clotting and centrifuged it for 5 min at 2000 xg. We collected the sera, stored it at −80°C, and used it for ELISA. We harvested BALF, lung, and spleen at the end of the study essentially as previously described^73,74^. We kept the sera and other biological fluids [with protease inhibitors (Roche, #11836153001] at -80 °C for long-term storage. After dissociation, the splenocytes and lung cells were frozen in FBS+10% DMSO and stored in the liquid nitrogen vapor phase until further use.

### ELISA

We determined the anti-N and anti-S antibody titers in serum or other biological fluids using ELISA. Briefly, we coated 0.5 μg/ml S protein (Acrobiosystems, DE, USA) and 1 μg/ml N protein (Sino Biological, PA, USA) onto ELISA plates (Corning, NY, USA) in PBS overnight at 4°C or for 2 h at 37 °C. The plate was then blocked with PBS+1% BSA (Fisher Scientific, PA, USA) +0.1% Tween 20™ (Sigma-Aldrich, MD, USA) for 2 h at RT. After washing, we added the samples at different dilutions. We detected the captured antibodies using HRP-conjugated anti-mouse IgG (Jackson ImmunoResearch Laboratories, 1 in 6,000; PA, USA), Goat anti-mouse IgA biotin (Southern Biotech, 1: 5,000; AL, USA). Streptavidin-HRP (Vector Laboratories, 1 in 2500, CA, USA) was used to detect the anti-IgA biotin antibodies. For BALF IgG ELISAs, antigens were coated onto plates at 0.5 μg/ml (S protein) and 2 μg/ml (N protein). We obtained the positive controls (anti-N and anti-S IgG) from Abeomics (CA, USA)

### Processing of spleen and lungs for ELISPOT and flow cytometry

To isolate lung cells, we perfused the lung vasculature with 5 ml of 1 mM EDTA in PBS without Ca^2+^, Mg^2+^ and injected it into the right cardiac ventricle. Each lung was cut into 100–300 mm^2^ pieces using a scalpel. We transferred the minced tissue to a tube containing 5 ml of digestion buffer containing collagenase D (2mg/ml, Roche #11088858001) and DNase (0.125 mg/ml, Sigma #DN25) in 5 ml of RPMI for 1 h and 30 min at 37 °C in the water bath by vortexing after every 10 min. We disrupted the remaining intact tissue by passage (6-8 times) through a 21-gauge needle. Next, we added 500 µL of ice cold-stopping Buffer (1x PBS, 0.1M EDTA) to stop the reaction. We then removed tissue fragments and dead cells with a 40 µm disposable cell strainer (Falcon) and collected the cells after centrifugation at 400 xg. We then lysed the red blood cells (RBCs) by resuspending the cell pellet in 3 ml of ACK Lysing Buffer (Invitrogen) and incubated for 3 min at RT, followed by centrifugation at 400 xg. Then, we discarded the supernatants and resuspended the cell pellets in 5 ml of complete RPMI medium (Corning, NY, USA). Next, we collected the spleen in RPMI medium and homogenized them through a 40 µm cell strainer using the hard end of a syringe plunger. After that, we incubated splenocytes in 3ml of ACK lysis buffer for 3 min at RT to remove RBCs, then passed through a 40 µm strainer to obtain a single-cell suspension. We counted the lung cells and splenocytes by the trypan blue exclusion method.

### ELISPOT

IFNγ and IL4 ELISpot assay was performed using Mouse IFNγ ELISPOT basic kit (ALP) and Mouse IL4 ELISPOT basic kit following the manufacturer’s instructions (Mabtech, VA, USA). For cell activation control, we treated the cultures with 10 ng/ml phorbol 12-myristate 13-acetate (PMA) (Sigma, St. Louis, MI, USA) and 1 µg/ml of ionomycin (Sigma, St. Louis, MI, USA). We used the complete medium (RPMI supplemented with 10% FBS) as the negative control. We stimulated splenocytes and lung cells (3 × 10^5^) *in vitro* with either N-protein peptide pool (Miltenyi Biotec; #130-126-699, Germany) or S-protein peptide pool (Genscript, # RP30020, USA) or S-protein (B.1.351) mutation peptide pool (Miltenyi Biotec, # 130-127-958, Germany) at a concentration of 1.5 μg/ml/peptide at 37 °C for 16-18 h in pre-coated ELISpot plate (MSIPS4W10 from Millipore) coated with AN18 IFNγ (1 µg/ml, Mabtech #3321-3-250;) and 11B11 IL4 (1 µg/ml, Mabtech #3311-3-250) coating antibody. The next day, we washed off the cells and developed the plates using biotinylated R4-6A2 anti-IFN-γ (Mabtech #3321-6-250) and BVD6-24G2 anti-IL4 (Mabtech #3311-6-250) detection antibody, respectively. Then, we washed the wells and treated them for 1 h at RT with 1:30,000 diluted Extravidin-ALP conjugate (Sigma, St. Louis, MI, USA). After washing, we developed the spots by adding 70 µL/well of BCIP/NBT-plus substrate (Mabtech #3650-10) to the wells. We incubated the plate for 20-30 min for color development and washed it with water. We quantified the spots using Cytation 7 (Bio-Tek Instruments, Inc.). Each spot corresponds to an individual cytokine-secreting cell. We showed the values as the background-subtracted average of measured triplicates.

### Cell surface staining, intracellular cytokine staining for flow cytometry

We stimulated the spleen and lung cells from immunized and control animals to detect nucleocapsid protein-specific CD8^+^ T cell responses with an N protein-peptide pool at a concentration of 1.5 μg/mL/peptide (Miltenyi Biotec; 130-126-699, Germany) at 37 °C for 16-18 h followed by the addition of Brefeldin A (5 μg/ml BD Biosciences #BD 555029) for the last 5 h of the incubation. We used 10 ng/ml PMA (Sigma, St. Louis, MI, USA) and 1 µg/ml ionomycin (Sigma, St. Louis, MI, USA) as the positive control. Stimulation without peptides served as background control. We collected the cells and stained with Live/Dead Aqua (Thermo Fisher #L34965) in PBS, followed by Fc-receptor blockade with anti-CD16/CD32 (Thermo Fisher #14-0161-85), and then stained for 30 min on ice with the following antibodies in flow cytometry staining buffer (FACS): anti-CD4 AF589 (clone GK1.5; Biolegend #100446), anti-CD8b (clone YTS156.7.7; Biolegend #126609), anti-CD69 (clone H1.2F3; Biolegend #104537), anti-CD137 (clone 1AH2; BD; # 40364), anti-CD45 (clone 30-F11; BD; #564279). We washed the cells twice with the FACS buffer. We then fixed them with 100 μL IC (intracellular) fixation buffer (eBioscience) for 30 min at RT. We permeabilized the cells for 10 min with 200 μL permeabilization buffer (BD Cytofix solution kit). We performed the intracellular staining using the antibodies Alexa Fluor 488 interferon (IFN) gamma (clone XMG1.2; BD; #557735) and Granzyme B (clone GB11; Biolegend; #515407) overnight at 4 °C. Next, we washed the cells with FACS buffer and analyzed them on LSR-Fortessa flow cytometer (BD Bioscience) using FlowJo™ software version 10.8 (Tree Star Inc, Ashland, OR, USA). We calculated the results as the total number of cytokine-positive cells with background subtracted. We optimized the amount of the antibodies by titration. See **Figure S10A** for the gating strategy.

### Viruses and biosafety

#### Viruses

Isolates of SARS-CoV-2 were obtained from BEI Resources (Manassas, VA) and amplified in Vero E6 cells to create working stocks of the virus. The virus was adapted to mice by four serial passages in the lungs of mice and plaque purified at Utah State University (USU).

#### Biosafety and Ethics

The animal experiments at USU were conducted in accordance with an approved protocol by the Institutional Animal Care and Use Committee of USU. The work was performed in the AAALAC-accredited LARC of the university in accordance with the National Institutes of Health Guide for the Care and Use of Laboratory Animals (8th edition; 2011).

### Viral challenge studies in animals

#### Animals

For SARS-CoV-2 animal studies completed at USU, 6 to 10-week-old male and female golden Syrian hamsters were purchased from Charles River Laboratories and housed in the ABSL-3 animal space within the LARC.

#### Infection of animals

Hamsters were anesthetized with isoflurane and infected by intranasal instillation of 1 × 10^4.5^ CCID_50_ (cell culture infectious dose 50%) of SARS-CoV-2 in a 100 µl volume.

#### Titration of tissue samples

Lung tissue and nasal tissue samples from hamsters were homogenized using a bead-mill homogenizer using minimum essential media. Homogenized tissue samples were serially diluted in a test medium, and the virus was quantified using an endpoint dilution assay on Vero E6 cells for SARS-CoV-2. A 50% cell culture infectious dose was determined using the Reed-Muench equation^75^.

### Histopathology

Lungs of the Syrian golden hamsters were fixed in 10% neural buffered formalin overnight and then processed for paraffin embedding. The 4-μm sections were stained with hematoxylin and eosin for histopathological examinations. A board-certified pathologist evaluated the sections.

### Quantification and statistical analysis

Statistical significance was assigned when P values were <0.05 using GraphPad Prism (v6.07). Tests, number of animals (n), mean values, and statistical comparison groups are indicated in the figure legends. Analysis of ELISA, ELISPOT, viral titers and pathology scores were performed using a Mann-Whitney test. We used a mixed-effects model for repeated measures analysis to compare bodyweight data at each time point.

### Quantitative modeling

To quantify the kinetics of SARS-CoV-2 infection in the upper respiratory tract (URT) upon N, S, and NS. immunization, we modified the innate immune model described by Ke et al. ^76^. We added appropriate parameters to account for de-novo blocking and T cell killing, as shown in **Figure 2A**. The mean population parameter values and initial values were taken from Ke et al. ^76^. We solved the system of ordinary differential equations (ODEs) for different S and N response efficiencies using the ODE45 function in MATLAB 2018b. A sample MATLAB code for solving the system of equations has been provided in supplementary methods.

## ACKNOWLEDGMENTS

This publication was supported by the NIH (R01GM143243), AuraVax Therapeutics, and Owens foundation. XL acknowledges partial funding support from the National Cancer Institute (NIH R15CA182769, P20CA221731, P20CA221696) and CPRIT (RP150656). The following reagent was produced under HHN272201400008C and obtained through BEI Resources, NIAID, NIH: Spike Glycoprotein (Stabilized) from SARS-Related Coronavirus 2, Wuhan-Hu-1, Recombinant from Baculovirus, NR-52308.

## AUTHOR CONTRIBUTIONS

NV conceived the study. NV, AL, AS, LJNC, MS, and BH designed the study. AL, AS, SRS, MK, MM, AD, RK, AR, SM, BH, and XL performed experiments. AL, AS, SRS, MK, BH, XL, and NV analyzed the data. MK performed modeling. NV and AL drafted the manuscript, and all authors contributed to the review and editing of the manuscript.

## DECLARATION OF INTERESTS

UH has filed a provisional patent based on the findings in this study. NV and LJNC are co-founders of AuraVax Therapeutics.

## SUPPLEMENTARY FIGURES

**Figure S1:**
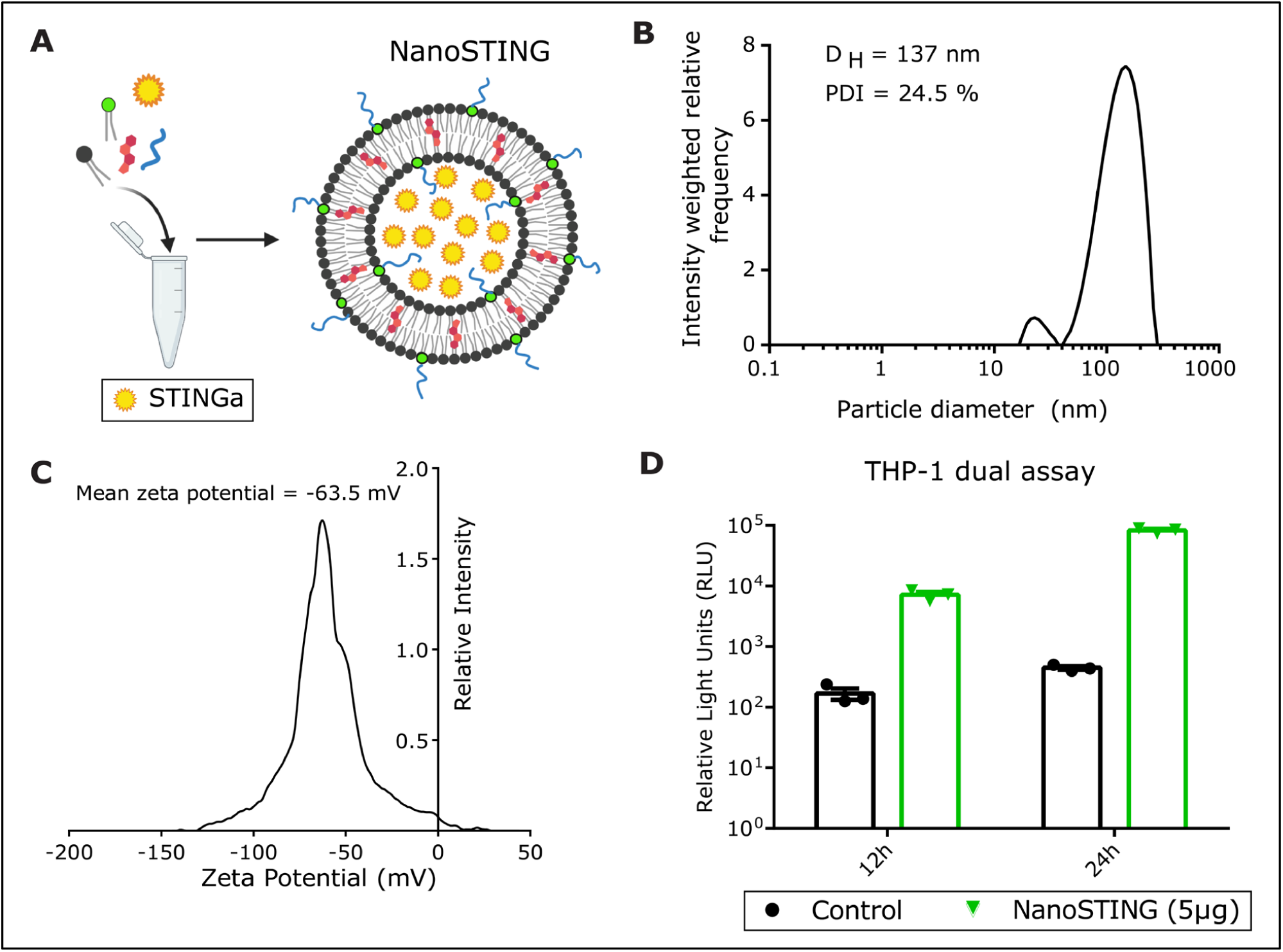
Synthesis and characterization of NanoSTING (related to Figure 1) **(A)** Overall schematic of the formulation of NanoSTING **(B)** Distribution of NanoSTING liposomal particle sizes measured by dynamic light scattering (DLS). D_H_: hydrodynamic diameter; PDI: polydispersity index **(C)** Zeta potential of the NanoSTING measured by electrophoretic light scattering (ELS). **(D)** Kinetics of the induction of luciferase in THP1-dual™ cells by NanoSTING (5 µg) at 12h and 24 h. RLU: relative light units. *Analysis was performed using a Mann-Whitney test. Vertical bars show mean values with error bar representing SEM. Mann-Whitney test: ****p < 0.0001; ***p < 0.001; **p < 0.01; *p < 0.05; ns: not significant*.

**Figure S2:**
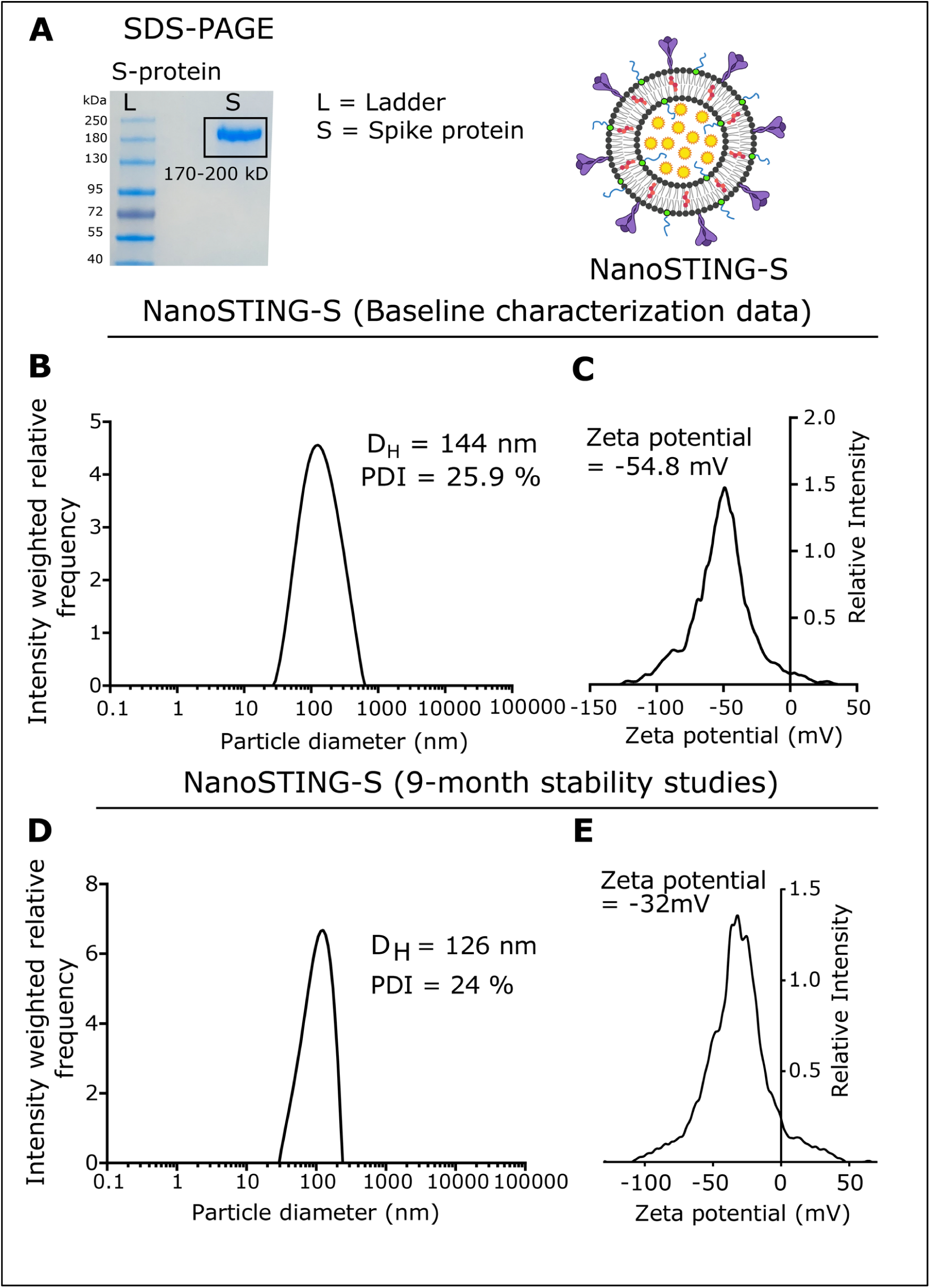
Characterization of NanoSTING-S (related to Figure 1) **(A)** Denaturing SDS-PAGE gel of the purified trimeric S protein. **(B)** Distribution of NanoSTING-S liposomal particle sizes measured DLS. **(C)** Zeta potential of the NanoSTING-S measured by ELS. **(D)** Distribution of NanoSTING-S particle sizes measured by DLS after storage at 4 °C for 9 months. **(E)** Zeta potential of the NanoSTING-S measured by ELS after storage 4 °C for 9 months.

**Figure S3:**
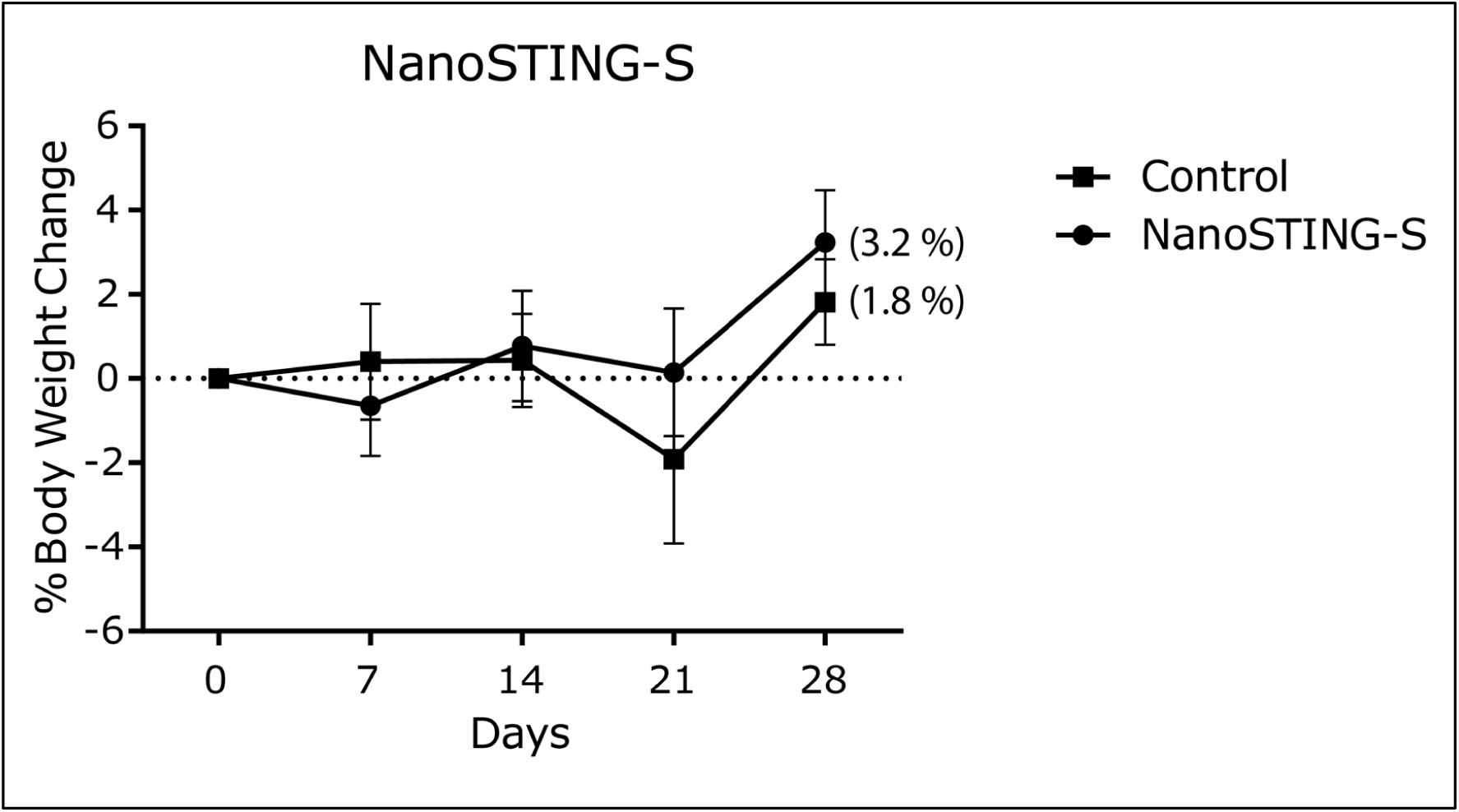
Percentage of bodyweight change of NanoSTING-S vaccinated mice compared to the control mice (related to Figure 1).

**Figure S4:**
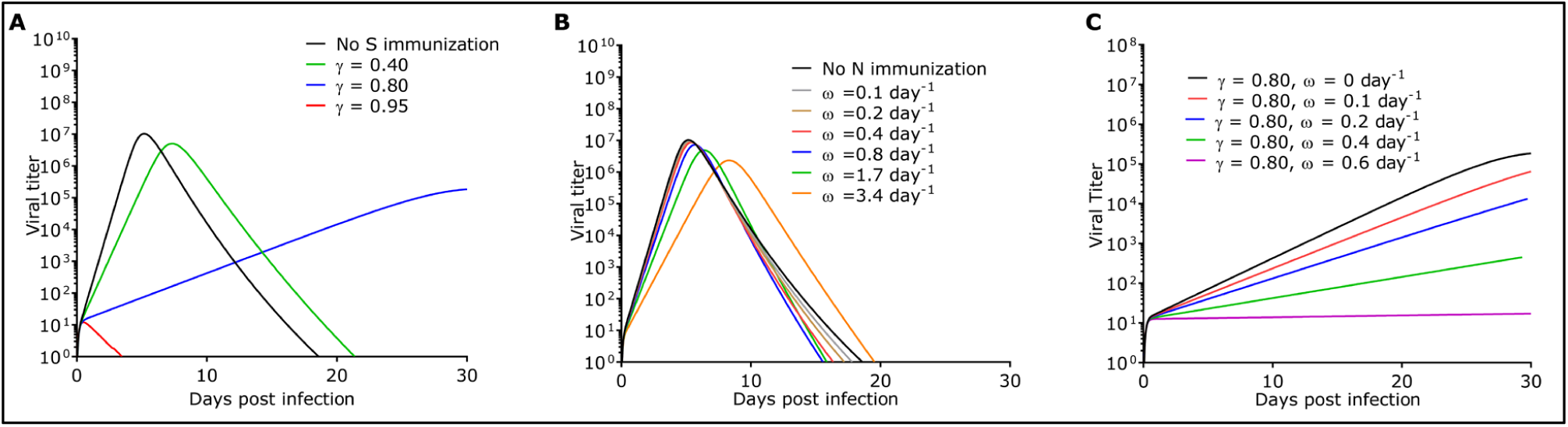
Evolution of viral dynamics with (A) S vaccine (assuming only de-novo blocking of viral entry) (B) N immunization (assuming only cytotoxic T cell killing of infected cells) and (D) vaccination with both N and S combined (related to Figure 2).

**Figure S5:**
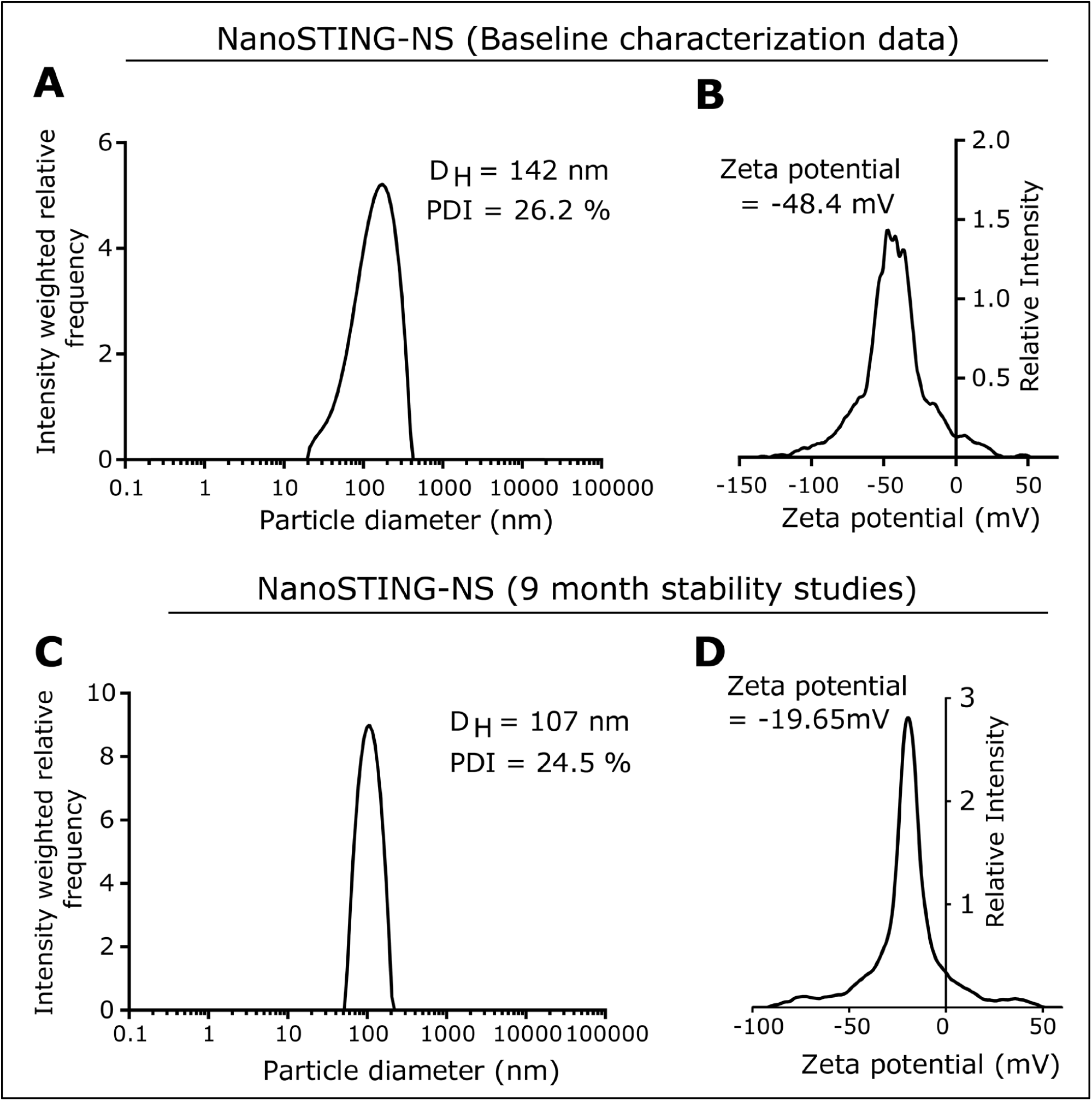
Characterization of NanoSTING-NS (related to Figure 3) **(A)** Distribution of NanoSTING-NS particle sizes measured by DLS. **(B)** Zeta potential of the NanoSTING-NS measured by ELS. **(C)** Distribution of NanoSTING-NS particle sizes measured by DLS after storage at 4 °C for 9 months. **(D)** Zeta potential of the NanoSTING-NS measured by ELS after storage 4 °C for 9 months.

**Figure S6:**
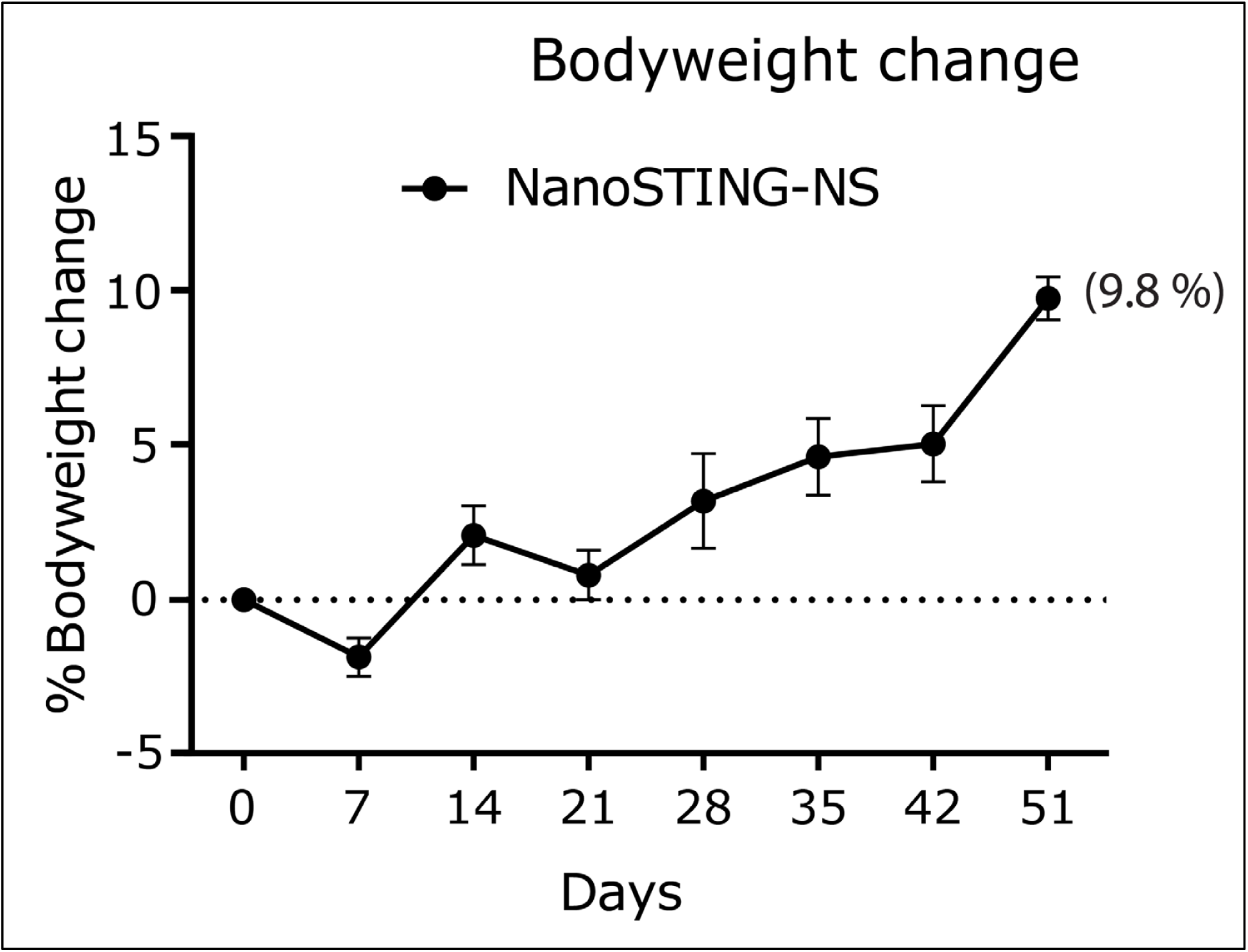
Percentage of bodyweight change of NanoSTING-NS vaccinated mice compared to the baseline (related to Figure 3).

**Figure S7:**
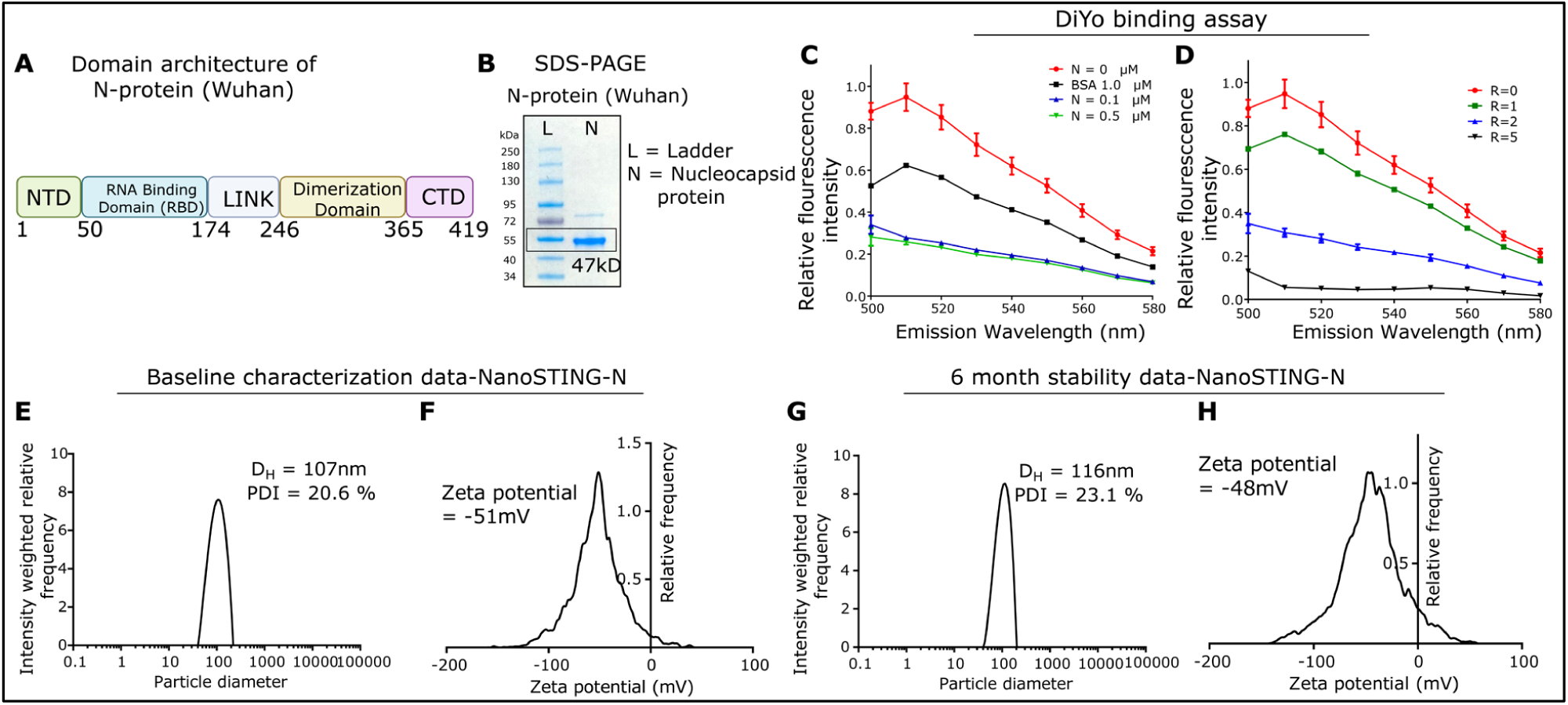
Characterization of NanoSTING-N (related to Figure 4). **(A)** Domain structure of the SARS-CoV-2 N protein. NTD: N-terminal domain; CTD: C-terminal domain^78^. **(B)** Denaturing SDS-PAGE gel of the purified nucleocapsid protein. **(C)** Fluorescence emission spectra of DNA-bound DiYO-1 in the presence of nucleocapsid-protein. Bovine serum albumin (BSA) was used as a control. **(D)** Fluorescence emission spectra of DNA-bound DiYO-1 in the presence of PEI (Positive control). **(E)** Distribution of NanoSTING-N particle sizes measured by DLS. **(F)** Zeta potential of the NanoSTING-N measured by ELS. **(G)** Distribution of NanoSTING-N particle sizes measured by DLS after storage at 4 °C for 6 months. (H) Zeta potential of the NanoSTING-N measured by ELS after storage at 4 °C for 6 months.

**Figure S8:**
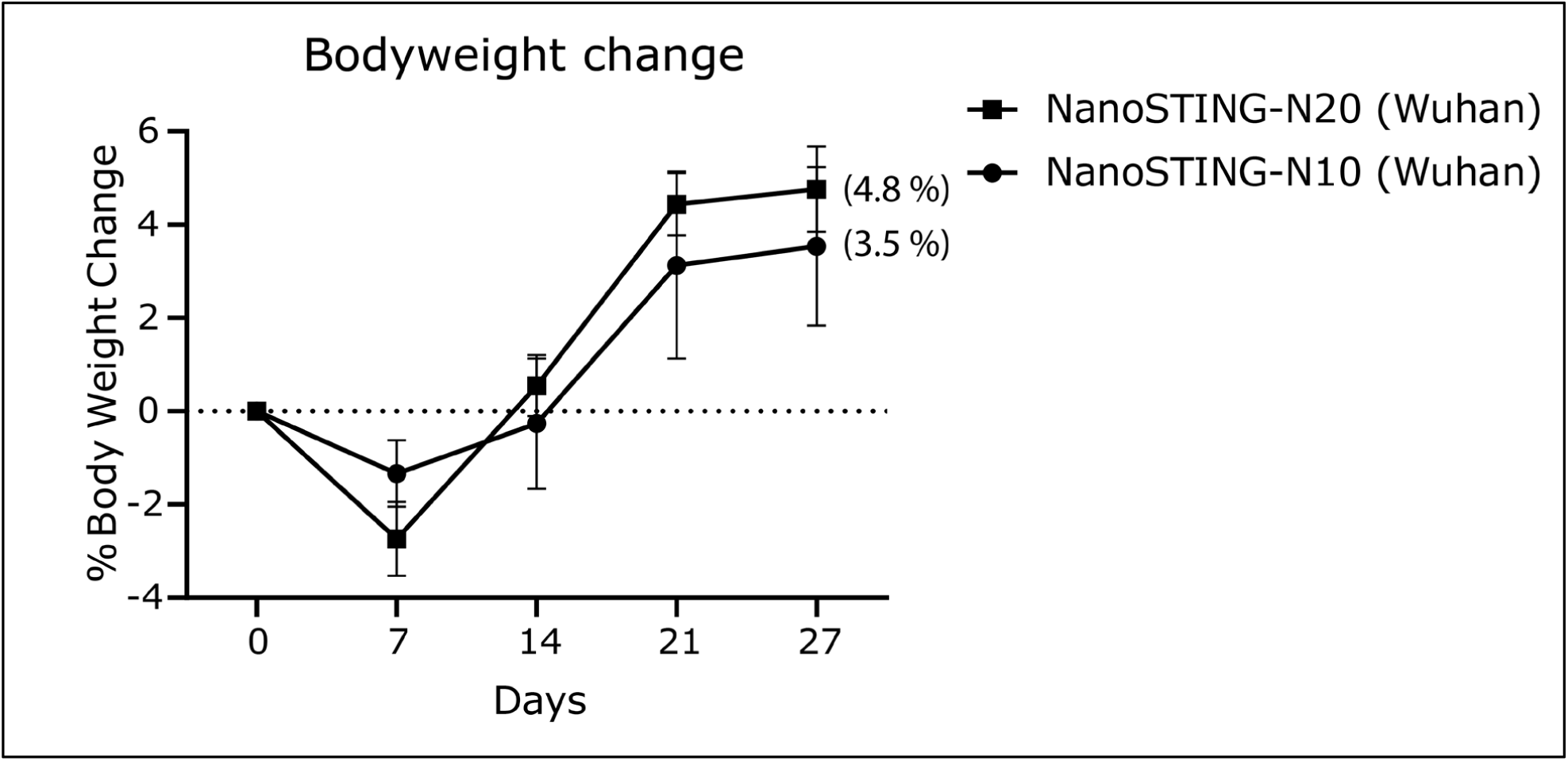
Percentage of bodyweight change of NanoSTING-N vaccinated mice compared to the baseline (related to Figure 4).

**Figure S9:**
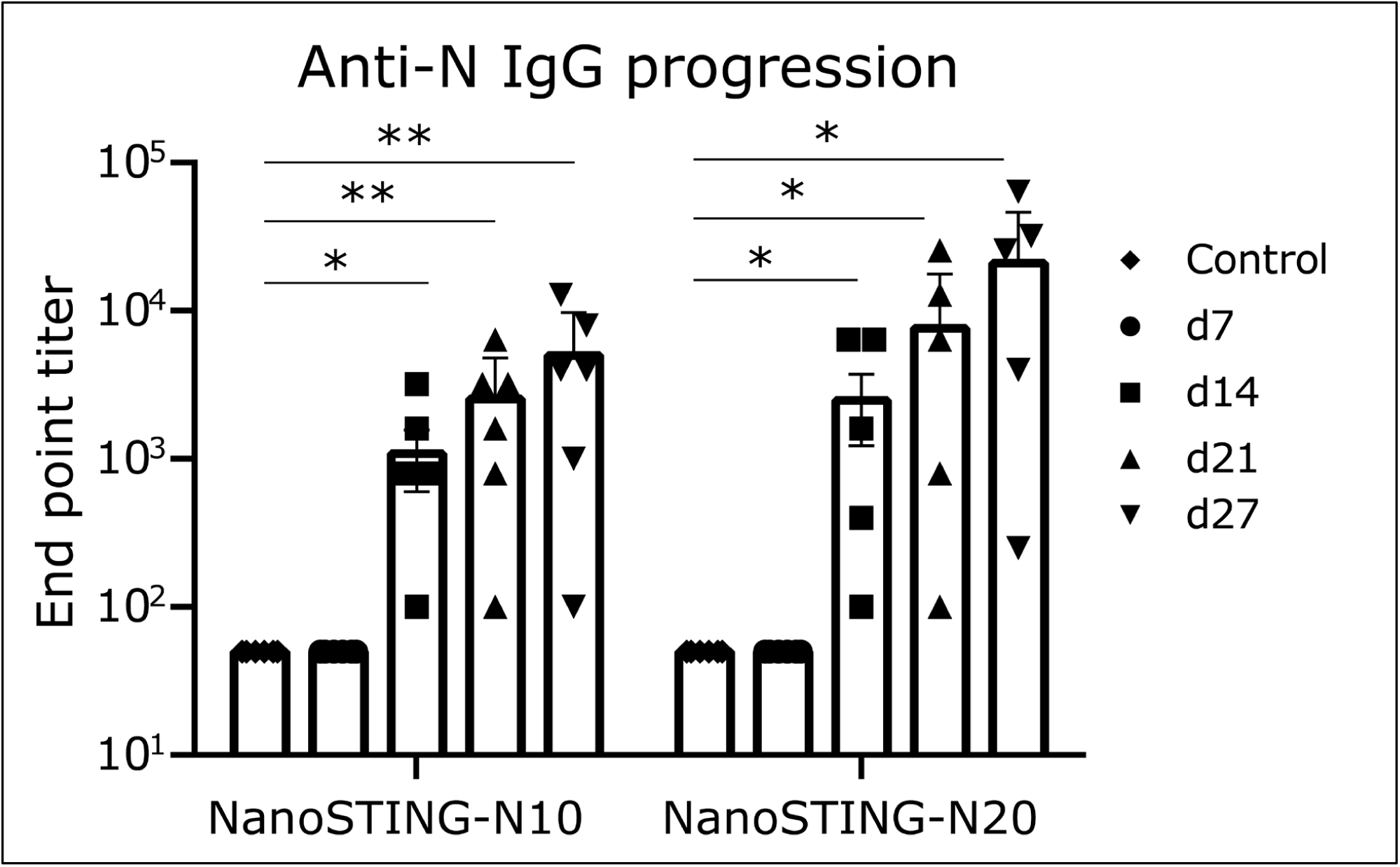
Anti-N IgG progression in the serum for NanoSTING-N10 and NanoSTING-N20 vaccinated mice at day 7, 14, 21, and 27 (related to Figure 4). *Analysis was performed using a Mann-Whitney test. Vertical bars show mean values with an error bar representing SEM. Each dot represents an individual mouse. Asterisks indicate significance compared to the non-vaccinated mice*. *\*\*\*\*p < 0.0001; ***p < 0.001; **p < 0.01; *p < 0.05; ns: not significant*.

**Figure S10:**
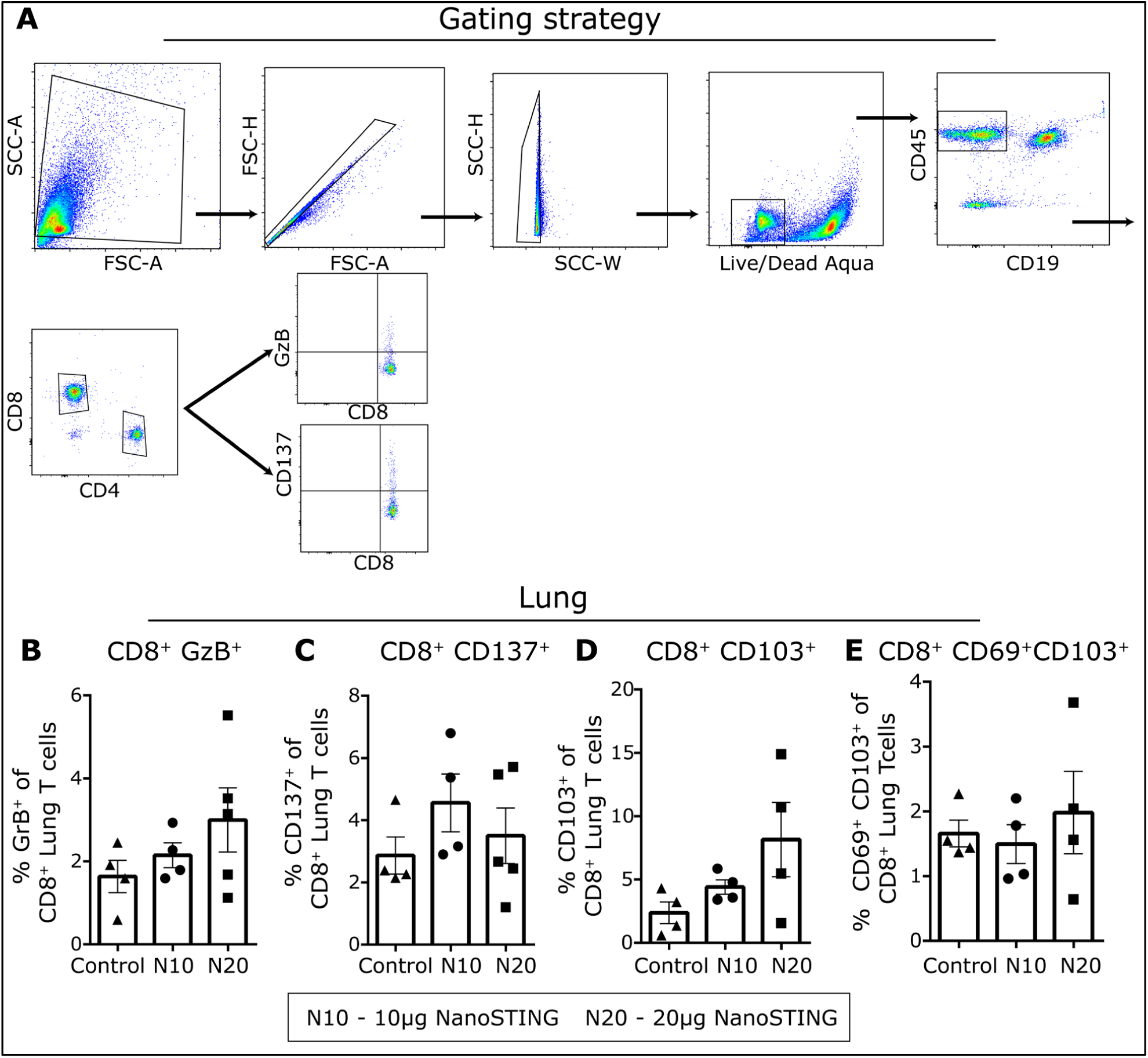
Lung CD8^+^ T cells were assayed for expression of Granzyme B (GrB), CD137, CD103, and CD69 by flow cytometry (related to Figure 4). **(A)** Flow cytometric gating strategy for the identification and quantification of N protein reactive CD8^+^ T cells **(B-E)** Lung cells were stimulated *ex vivo* with overlapping peptide pools, and the expression of GzB^+^ (B), CD137^+^ (C), CD103^+^ (D), CD103^+^ and CD69^+^ (E) expression in CD8^+^ T cells were quantified using flow cytometry. *Analysis was performed using a Mann-Whitney test. Vertical bars show mean values with error bar representing SEM. Mann-Whitney test: ****p < 0.0001; ***p < 0.001; **p < 0.01; *p < 0.05; ns: not significant*.

**Figure S11:**
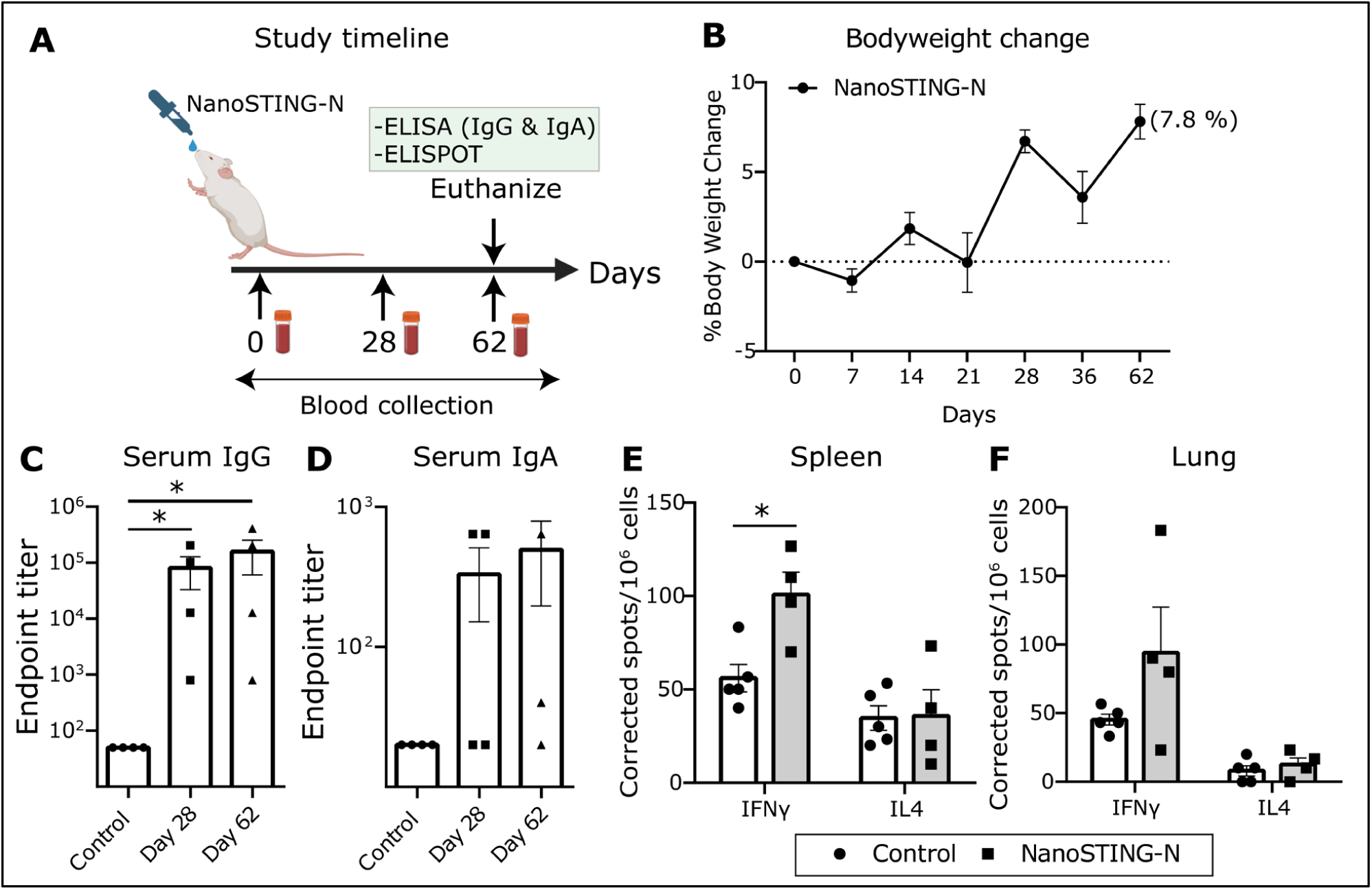
Immunization with NanoSTING-N elicits strong cellular and humoral immune responses (related to Figure 4) **(A)** Experimental set up: We immunized Balb/c mice (n=4/group) intranasally with single dose of NanoSTING-N followed by collection of serum after day 28 and day 62. The bodyweights of the animals were monitored every week after the immunization. We euthanized the animals at day 62 followed by the collection of serum, lungs, and spleen. Bodyweight change, ELISA (IgG & IgA), and ELISPOT (IFNγ and IL4) were used as primary endpoints. Naïve Balb/c mice were used controls (n=4-5/group). **(B)** Percent bodyweight change compared to the baseline at the indicated time intervals. **(C, D)** Humoral immune responses in the serum were evaluated using N-protein based IgG and IgA ELISA. **(E, F)** Splenocytes (E) and lung cells (F) were stimulated *ex vivo* with overlapping N and S-peptide pools, and IFNγ & IL4 responses were detected by ELISPOT. *For ELISA and ELISPOT data, analysis was performed using a Mann-Whitney test. Vertical bars show mean values with error bar representing SEM. Each dot represents an individual mouse*. *\*\*\*\*p < 0.0001; ***p < 0.001; **p < 0.01; *p < 0.05; ns: not significant*.

## SUPPLEMENTARY METHODS

To quantify the kinetics of SARS-CoV-2 infection in the upper respiratory tract (URT) upon different antigens used for immunization (Spike(S), Nucleocapsid(N) and N+S), we modified the innate immune model as described previously^76^. If S immunization works by *de novo* blocking of viral entry through neutralizing antibodies and N immunization works by inducing cytotoxic T cell responses, we modified the governing equations as shown in the **Table1** below and **Figure 2** of the main manuscript.

**Table 1:** Equations governing anti-viral response for different modes of immunizations. T = Target cells, R = Refractory cells, E = Eclipse phase cell (Infected cells not producing virus), I = Infected cells productively making virus, V = Viral load

| No immunization | S immunization | N immunization | N+S immunization |
| --- | --- | --- | --- |
| $\frac{dT}{dt} = -\beta VT - \phi IT + \rho R$<br>$\frac{dR}{dt} = \phi IT - \rho R$<br>$\frac{dE}{dt} = \beta VT - kE$<br>$\frac{dI}{dt} = kE - \sigma I$<br>$\frac{dV}{dt} = \pi I - cV$ | $\frac{dT}{dt} = -\beta(1-\gamma)VT - (\phi I)T + \rho R$<br>$\frac{dR}{dt} = (\phi I)T - \rho R$<br>$\frac{dE}{dt} = \beta(1-\gamma)VT - kE$<br>$\frac{dI}{dt} = kE - \sigma I$<br>$\frac{dV}{dt} = \pi I - cV$ | $\frac{dT}{dt} = -\beta VT - \phi IT + \rho R$<br>$\frac{dR}{dt} = \phi IT - \rho R$<br>$\frac{dE}{dt} = \beta VT - kE$<br>$\frac{dI}{dt} = kE - (\sigma + \omega)I$<br>$\frac{dV}{dt} = \pi I - cV$ | $\frac{dT}{dt} = -\beta(1-\gamma)VT - (\phi I)T + \rho R$<br>$\frac{dR}{dt} = (\phi I)T - \rho R$<br>$\frac{dE}{dt} = \beta(1-\gamma)VT - kE$<br>$\frac{dI}{dt} = kE - (\sigma + \omega)I$<br>$\frac{dV}{dt} = \pi I - cV$ |

We solved these ordinary differential equations with mean population parameter values and initial values taken from Ke et al and as shown in **Table 2 & 3**. We calculated the Viral load area under the curve (AUC) during infection for varying de-novo blocking efficiency (γ) and cytotoxic killing efficacy (ω).

**Table 2:** Mean population values of parameters for equations in Table 1.

| Parameter | Description | Mean population value/unit |
| --- | --- | --- |
| $\beta$ | Infectivity parameter constant | $3.2 \times 10^{-8}$ ml/RNA copy/day |
| $\sigma$ | Death rate of infected cells | 1.7 /day |
| $\pi$ | Composite parameter for virus production and sampling | 45.3/ml/day |
| $\phi$ | Rate constant for interferon induced conversion of Target cells to refractory cells | $1.3 \times 10^{-6}$ /cell/day |
| k | 1/the eclipse phase duration | 4 /day |
| c | virus clearance rate | 10 /day |
| $\rho$ | Rate at which refractory cells become target cells again | 0.0044/day |
| $\gamma$ | De-novo blocking efficacy (Antibody efficiency in blocking viral entry) | 0-1 (Variable) |
| $\omega$ | Death rate of infected cells mediated by cytotoxic T cells | Variable /day |

**Table 3:**
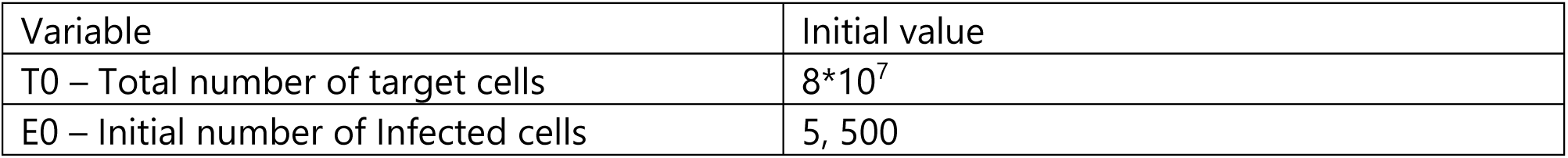
Initial values for total number of target cells(T0) and initial number of infected cells(I0) upon viral exposure.

Sample MATLAB code:

~~~
% Covid dynamics with NanoSTING
%beta - Infectivity parameter constant = 3.2*10^-8 ml/RNA copy/day
%delta - Death rate of infected cells - 1.7/day
%pii - Composite parameter for virus production and sampling - 45.3/ml/day
%phi - Rate constant for interferon induced conversion of Target cells to
%refractory cells - 1.3*10^-6 /cell/day
%rho - Rate at which refractory cells become target cells again -0.0044/day
%c - Virus clearance rate - 10/day
%k - 1/the eclipse phase duration = 4/day
~~~

~~~
beta = 3.2*10^-8; %ml/RNA copy/day
delta = 1.7; %/day
pii = 45.3; %/ml/day
phi = 1.3*10^-6; %/cell/day
rho = 0.0044; %/day
c = 10; %/day
k = 4; %/day
denovo = 0.8; % coefficient responsible for denovo-blocking
cyT = 0.6; % % killing rate of cytotoxic T cells
T0 = 8*10^7; %Total number of target cells
R0 = 0; %Initial refractory cells
E0 = 5; %Initial number of infected cells
I0 = 0;
V0 = 0; %Initial virus titer
~~~

~~~
t_int = [0,30];
init_cond = [T0,R0,E0,I0,V0]’;
[t,y] = ode45(@(t,Y) covidode(t,Y,beta,delta,pii,phi,rho,c,k,denovo,cyT), t_int,init_cond);
-----------------------------------------------------------------------------
~~~

Function handle:

~~~
function dYdt = covidode(t,Y,beta,delta,pii,phi,rho,c,k,denovo,cyT)
dYdt = [-beta*(1-denovo)*Y(5)*Y(1)-(phi*Y(4))*Y(1)+rho*Y(2);
    (phi*Y(4))*Y(1)-rho*Y(2);
    beta*(1-denovo)*Y(5)*Y(1)-k*Y(3);
    k*Y(3)-delta*Y(4)-cyT*Y(4);
    pii*Y(4)-c*Y(5)];
end
~~~

